# BdNRT2A and BdNRT3.2 are the major components of the High-Affinity nitrate Transport System in *Brachypodium distachyon*

**DOI:** 10.1101/2023.11.24.567652

**Authors:** Laure C. David, Mathilde Grégoire, Patrick Berquin, Anne Marmagne, Marion Dalmais, Abdelhafid Bendahmane, Anthony J. Miller, Anne Krapp, Françoise Daniel-Vedele, Thomas Girin, Sylvie Ferrario-Méry

## Abstract

- An efficient nitrate uptake system contributes to the improvement of crop nitrogen use efficiency under low nitrogen availability. The <u>H</u>igh <u>A</u>ffinity nitrate <u>T</u>ransport <u>S</u>ystem (HATS) in plants is active in low external nitrate and is mediated by a two-component system [high affinity transporters NRT2 associated to a partner protein NRT3 (NAR2)].
- In Brachypodium, the model plant for C3 cereals, we investigated the role of *BdNRT2A* and *BdNRT3.2* through various experimental approaches including gene expression profiling, functional characterisation in heterologous system, intracellular localization by imaging, and reverse genetics via gene silencing.
- Expression of *BdNRT2.A* and *BdNRT3.2* genes in response to nitrate availability fits with the characteristics of the HATS components. Co-expression of *BdNRT2A* and *BdNRT3.2* is required for an effective nitrate transport in the heterologous expression system Xenopus oocytes. Functional interaction between BdNRT2A-GFP and BdNRT3.2-RFP fusion proteins has been observed at the plasma membrane in Arabidopsis protoplasts in transient expression experiments. BdNRT3.2 appeared to be necessary for the plasma membrane localization of BdNRT2A. ^15^Nitrate influx measurements with *bdnrt2a* mutants (two amiRNA mutants and one NaN_3_ induced mutant with a truncated NRT2A protein), confirmed that BdNRT2A is a major contributor of the HATS in Brachypodium.
- Directed mutagenesis in BdNRT2A of a conserved Ser residue (S461) specific to monocotyledons has been performed to mimic a non-phosphorylated S461A or a constitutively phosphorylated S461D, in order to evaluate its potential role in the BdNRT2A and BdNRT3.2 interaction leading to plasma membrane targeting. Interestingly, the phosphorylation status of S461 did not modify the interaction, suggesting on a more complex mechanism.
- In conclusion, our data show that BdNRT2A and BdNRT3.2 are the main components of the nitrate HATS activity in Brachypodium (Bd21-3) and allow an optimal growth in low N conditions.

## Introduction

Nitrogen (N) is an essential nutrient for the growth and development of plant, and is present in the soil in the form of nitrate (NO_3_^-^) and ammonium (NH_4_^+^). To ensure high crop yields, N - containing chemicals fertilizers have largely been used for 60-70 years to provide NO_3_^-^ and NH_4_^+^ to plants and thus to ensure enough food to the constantly increasing human population. Attempts to improve the Nitrogen Use Efficiency (NUE) of crops are considered as a priority goal for the near future in order to limit the fertilizer over-use and the related environmental problems, such as eutrophication. It is assumed that NUE can be improved by enhancing NO_3_^-^ uptake capacity, the major N source for most plants in aerobic conditions (Bogard *et al*., 2010; Chen *et al*., 2016; Wang *et al*., 2018). Depending on the external NO_3_^-^ concentrations, two different uptake systems occur within the plant: the high-affinity (HATS) and low-affinity (LATS) systems operate when NO_3_^-^ is present in the soil at low (< 1 mM) or high concentration (> 1 mM), respectively. NO_3_^-^ is taken up by roots from the soil by members of NRT2 (Nitrate Transporter 2) and NPF (Nitrate Transporter 1/Peptide Transporter Family) families. NRT2 proteins are active when the soil NO_3_^-^ concentration is low (below 1mM), thus contributing to the HATS for NO_3_^-^ uptake. Molecular characterization of NRT2 genes have been largely performed in Arabidopsis. Among the seven genes encoding NRT2 transporters (*AtNRT2.1* to *2.7*) (Wang *et al*., 2018), four have been characterized *in planta* as involved in root NO_3_^-^ uptake, AtNRT2.1 being more active than AtNRT2.2 (Filleur *et al*., 2001; Li *et al*., 2007) and AtNRT2.4 and AtNRT2.5 being effective at very low NO_3_^-^ concentrations (Kiba *et al*., 2012; Lezhneva *et al*., 2014). Two other AtNRT2 have functions in aerial parts of the plants: AtNRT2.6 is involved in biotic interaction in leaves (Dechorgnat *et al.,* 2012) and AtNRT2.7 is responsible for vacuolar NO_3_^-^ loading in seeds (Chopin *et al.,20*07).

Plants modulate root NO_3_^-^ uptake by regulating transcript abundance and protein activity, depending on their carbon and N status. Thereby, expression of *AtNRT2.1* is stimulated by low NO_3_^-^ concentration, light and sugars (Filleur and Daniel-Vedele, 1999; Lejay *et al*., 1999; Okamoto *et al*., 2003) and repressed by NH_4_^+^ and amino acids (Lejay *et al*., 1999; Zhuo *et al*., 1999; Nazoa *et al*., 2003). Nevertheless, studies of plants overexpressing *AtNRT2.1* reveal that despite a constitutively high expression, HATs activity can be repressed (Laugier *et al*., 2012) indicating that a post translational regulation interferes, leading to an inactivation of the transporter. More recently, post-translational regulation of AtNRT2.1 have been identified at the N and C-terminal ends. Phosphorylation of Ser28 in the N-terminus of AtNRT2.1 results in the protein stabilisation in response to NO_3_^-^ limitation (Zou *et al*., 2020) and phosphorylation of Ser501 inactivates NRT2.1 and thus NO_3_^-^ transport activity in response to NH_4_^+^ supply (Jacquot *et al*. 2020). Some NRT2s were shown to require a partner protein NRT3 (also known as NAR2) for function (Tong *et al*., 2005).

Proteomic studies in Arabidopsis have revealed the functional structure of the high-affinity NO_3_^-^ transport, composed of two AtNRT2.1 and two AtNRT3.1 (NAR2) subunits (Yong *et al*., 2010). Two *AtNRT3* genes have been identified but *AtNRT3.1* expression is predominating and is induced by NO_3_^-^ (Okamoto *et al*. 2006; Orsel *et al*., 2006). Analysis of *atnrt3.1* mutants revealed that the absence of AtNRT3.1 protein affects the NO_3_^-^-inducible component of HATS (Okamoto *et al*. 2006) and the localization of AtNRT2.1 at the plasma membrane (pm) (Wirth *et al*., 2007). The molecular mechanism of NRT2 and NRT3 interaction has started to be elucidated. NRT2 protein structure is predicted to contain 12 transmembrane domains and N and C-terminal ends directed towards the cytosol (Jacquot *et al*. 2020) and the NRT3 protein contains only one transmembrane domain (Tong *et al*. 2005). The AtNRT2.1/AtNRT3.1 association is related to the N-terminal, the C-terminal ends of AtNRT2.1, and to the central part of NRT3. Leu85 located in the first transmembrane domain of AtNRT2.1 is critical for the association between AtNRT2.1 and AtNRT3.1 (Kotur *et al*., 2017). In addition to Arabidopsis, capacity of transporting NO_3_^-^ through the interaction with NRT3 protein have been proved in various dicotyledonous and also monocotyledonous species, as for HvNRT2.1 (Tong *et al*., 2005), and OsNRT2.1 and OsNRT2.3a (Yan *et al*., 2011). In rice 30 amino acids (from 65 to 95) at the N-terminal end of OsNRT2.3a are required for the interaction with OsNAR2.1 and for the NO_3_^-^ transport activity (Feng *et al*., 2011). The central region of NRT3 has been indicated to interact with NRT2 in rice, where two residues (R100 and D109) placed in the middle region of OsNAR2.1 are necessary for the interaction with OsNRT2.3a at the pm (Liu *et al*., 2014). In Arabidopsis the replacement of D105 in AtNRT3.1 markedly reduced NO_3_^-^ uptake (Kawachi *et al*., 2006). But the C-terminus region of NRT2 has been also indicated to interact with central region of NRT3 in barley, where the Ser463 of HvNRT2.1 was shown to control the interaction between HvNRT2.1 and HvNAR2.1, and it was suggested that the interaction is regulated by the phosphorylation/dephophorylation of Ser463 (Ishikawa *et al*., 2009). The phosphorylation site Ser501 at the C-terminus of AtNRT2.1 was shown to inactivate the transporter leading to a decrease in HATS activity, but clearly independently of the breaking of AtNRT2.1 and AtNRT3.1 interaction (Jacquot *et al*., 2020). Intriguingly, this Ser is replaced by a Gly in monocotyledons suggesting distinct regulation mechanisms between monocotyledons and dicotyledons. (Jacquot *et al*., 2017).

NRT2.1 orthologs have been identified in numerous species as rice (Cai *et al*., 2008), barley (Vidmar *et al*., 2000), wheat (Yin *et al*., 2007), tomato (Ono and Frommer, 2000), tobacco (Alberto *et al*., 1997), rapeseed (Faure-Rabasse *et al*., 2002) and peach (Nakamura *et al*., 2007). In *Brachypodium distachyon*, the model plant for C3 cereals, we have previously identified seven NRT2 and two NRT3 (Girin *et al*., 2014). In the phylogenetic tree of the *NRT2* family, we identified five Brachypodium genes that cluster together in the same clade as AtNRT2.1 and HvNRT2.1 (Girin *et al*., 2014). It was impossible to predict which one was a functional orthologous of AtNRT2.1 among the five BdNRT2. We previously observed that the HATS activity was regulated by N availability and correlated with the expression of two *BdNRT2* (*BdNRT2A/2B*) and one *BdNRT3* (*BdNRT3.1*) genes, suggesting these BdNRTs are good candidates for elements of HATS activity (David *et al*., 2019). In this study, we demonstrated the major role of BdNRT2A/BdNRT3.1 in the HATS activity by using *bdnrt2a* mutants. We also studied the role of a Ser residue (S461) conserved in monocotyledons, but not in dicotyledons, for the interaction between BdNRT2A and BdNRT3.2.

## Materials and Methods

### Plant material and culture conditions

*Brachypodium distachyon* accession Bd21-3 was used for all experiments.

For the NO_3_^-^ influx measurements, Brachypodium seeds were germinated for four days in water at room temperature conditions. Seedlings plants were then transferred to hydroponic conditions in a growth chamber (18 h light at 22° C /6 h dark at 18°C cycle and 250 μmol photons m^-^². s^-1^ irradiation, OSRAM Lumilux L36W865 cool day light). Plants were provided with media containing 0.2 mM NO_3_^-^ : 0.05 mM Ca(NO_3_)_2_, 0.1 mM KNO_3_, 1 mM KH_2_PO_4_, 3.25 mM CaCl_2_, 5.45 mM KCl, 2 mM MgSO_4_, 4.5μM MnCl_2_,10 μM H_3_BO_3_, 0.7 μM ZnCl_2_, 0.4 μM CuSO_4_, 0.22 μM MoO_4_Na_2_, 50 μM iron–EDTA. For 1 mM and 0.1mM of nitrate, media contain 0.5 mM Ca(NO_3_)_2_ or 0.05 mM Ca(NO_3_)_2_ respectively and potassium was compensated with 5 mM and 5.45 mM of KCl respectively. For the nutrient solution containing 0.02 mM NO_3_^-^, nitrate was supplemented with 0.01 mM KNO_3_ and 0.005 mM Ca(NO_3_)_2_ and with 5 mM KNO_3_ and 2.5 mM Ca(NO_3_)_2_ for the 10 mM NO_3_^-^ nutrient solution.

For the NO_3_^-^ induction experiments, Brachypodium plants were grown in hydroponics, as described above, for 18 days on 0.1 mM NO_3_^-^, then they were NO_3_^-^ starved for 4 days (KCl was supplemented instead of KNO_3_) and finally re-supplied with 1 mM NO_3_^-^ for 2h or 3 h before harvest.

For the complementation study and protoplasts transfection study, WT and mutant seeds of Arabidopsis, ecoptype Wassilewskija (Ws) (*A. thaliana*), were used. For the complementation study plants were grown hydroponically in a growth chamber with 8 h light at 21° C /16h dark at 17°C cycle, 80% relative humidity and 150 μmol photons m ^-^² s^-1^ irradiation. Seeds were sterilized and stratified in water at 4 C for 5 days before sowing. Each seed was sown on top of a cut Eppendorf tube filled with medium consisting of half-strength nutrient solution containing 0.8% agar. Plants were supplied with media containing 0.2 mM NO_3_^-^ : 0.1 mM KNO_3_, 0.05 mM Ca(NO_3_)_2_ and 2.45 mM K2SO4, 2.15 mM CaCl_2_, 2 mM MgSO_4_, 2 mM KH_2_PO_4_,10 μM MnSO_4_, 24 μM H_3_BO_3_, 3μM ZnSO_4_, 0.9 μM CuSO_4_, 0.04 μM(NH_4_)_6_Mo_7_O_24_, iron–EDTA 10 mg l^-1^. The nutrient solution was changed every 3 days and, during the first 2 weeks was used at half-strength media. Plants were harvested 40 days after sowing, 1 h after illumination had started and analyzed for nitrate influx. Shoots and roots were weighed separately and frozen in liquid nitrogen. Influxes of ^15^NO_3_^-^ were performed on wild type genotype (Ws), mutant (*atnrt2.1-1*) and *atnrt2.1-1* mutant lines overexpressing *Pro35S::BdNRT2A* or *Pro35S::BdNRT2A*-GFP or *Pro35S::GFP-BdNRT2A* after 42 days of hydroponic growth.

For the protoplast transfection study Arabidopsis seeds were surface sterilized and sown on *in vitro* plates containing 1 % agar (Sigma-Aldrich France) as previously described (David *et al.,* 2016) and supplemented with a 9 mM NO_3_^-^ medium : 2 mM Ca(NO_3_)_2_, 5 mM KNO_3_, 2.5 mM KH_2_-PO_4_, 2 mM MgSO_4_, 0.07 % MES (pH 6), 0.005 % (NH_4_)_5_Fe(C_6_H_4_O_7_)_2_, 70µM H_3_BO_3_, 14µM MnCl_2_, 0.5 µM CuSO_4_, 10 µM NaCl, 1 µM ZnSO_4_, 0.001 µM CoCl_2_, and 0.2 µM NH_4_MoO_4_. After 3 days of stratification at 4 °C in the dark, plates were placed in a growth chamber at 18 °C with a 16/8 h light photoperiod, 60 % of humidity, and a light intensity of 50 µmol photons m^-2^ s^-1^. Seedlings were harvested 2 weeks after the transfer in light conditions for protoplast purification and transfection.

### ^15^NO_3_^-^ uptake in heterologous expression system (*Xenopus laevis* oocytes)

The coding sequences of BdNRT2A and BdNRT3.2 were cloned into pGEMT easy vector (Promega) and then digested with NotI enzyme. cDNA fragments were blunted using the Klenow fragment and subcloned in the EcoRV site of the pT7TS expression vector containing the 5’-untranslated region (UTR) and 3’-UTR of the Xenopus β-globin gene (Cleaver *et al.,* 1996). Clones with correct sequence were used for *in vitro* synthesis of RNA, pT7TS clones were linearized by digestion with XbaI. Capped full-length cRNAs were synthesized using a T7 RNA transcription kit (mMESSAGE mMACHINE; Ambion). As described in Orsel *et al*., (2006), we used a heterologous expression system *Xenopus laevis* oocytes. Xenopus oocytes were prepared as described previously (Zhou *et al*., 1998) and stored in ND96 solution (96 mM NaCl, 2 mM KCl, 1.80 mM CaCl_2_,1mM MgCl_2_, 15 mM MES, adjusted at pH 6 with NaOH). Healthy oocytes at stage V or VI were injected with 50 nL of water (nuclease free) or different cRNAs at 1mg/mL each. After 3-days incubation at 18°C, five to 10 oocytes were incubated in 3 mL of ND96 solution enriched with 0.5 mM Na^15^NO_3_ (atom%^15^N: 98%) during 16 h at 18°C. The oocytes were then thoroughly washed four times with ice-cooled 0.5 mM NaNO_3_ ND96 solution and dried at 60°C. The ^15^N to ^14^N ratio of single dried oocyte was measured using an isotope ratio mass spectrometer (model Integra CN; PDZ Europa). The delta ^15^N was calculated as described previously (Tong *et al.,* 2005).

### Identification of chemical mutagenesized mutants with mutations in the coding sequence of *BdNRT2A* in the TILLING collection of Versailles

The NaN_3_-induced mutant collection from Versailles (Dalmais *et al*., 2013) was used to search point mutations by a TILLING method in the coding sequence of *BdNRT2A.* The genomic DNA pools corresponding to 5530 M2 families were screened by PCR using *BdNRT2A* specific primers fused to fluorochromes. Mutations were then identified by sequencing the PCR products after digestion by restriction endonuclease ENDO1 and electrophoresis detection by laser of the cleaved amplicons. The effect of each point mutation was analysed and predicted using SIFT (Sorting Intolerant From Tolerant) program (http://sift.jcvi.org/). A SIFT score lower than 0.05 predicted a deleterious amino acid substitution for a point mutation.

### *BdNRT2A* amiRNA mutants

The amiRNA constructs were engineered using the online microRNA designer WMD3 (http://wmd3.weigelworld.org/cgi-bin/webapp.cgi). Specific sequences were designed to target *BdNRT2A* (TAAAGACAGCAGCAGTCGCGG) or the five *NRT2* genes BdNRT2A/B/C/D/F (TATCATGATGCGCACCTACTA). DNA fragments containing the specific sequences, the microRNA structure of pNW55 (based on the rice osa-MIR528; Warthmann *et al*., 2008) and Gateway attL1/attL2 borders were synthetized and cloned at the EcoRV site of pUC57-Kan (GenScript Biotech Corporation) (Supplementary data S1). These entry clones were subsequently recombined into the destination vector pIPKb002 (Himmelbach *et al*., 2007) using the Gateway LR Clonase II enzyme (Invitrogen). This vector contains the Hpt plant selection marker (hygromycin resistance) and drives the expression of amiRNAs under the constitutive *ZmUbi1* promoter. Transgenes were integrated into the B. distachyon accession Bd21-3 genome by Agrobacterium-mediated transformation (Agrobacterium tumefaciens strain AGL1) of embryogenic calli following the protocol described by Vogel and Hill (2008). Homozygous lines were selected based by hygromycin resistance.

### GFP fusions and functional complementation *of atnrt2.1-1* with *BdNRT2A*

The cDNA of *BdNRT2A* was first amplified by PCR using specific forward primer *70F2* and reverse primer *70R2* before being cloned into the pGEMT-easy vector (Promega). Clones with correct sequence were used to produced appropriate PCR product for Gateway cloning. First primers Bd70start and Bd70end or Bd70stop were used and PCR products were amplified with the universal *U3endstop* and *U5* primers to create the recombinant sites AttB. The product of recombination reactions (BP reactions) was used to transform competent *Escherichia coli* strain TOP10 (Invitrogen), by heat shock. LR clonase reactions to transfer fragments from the entry clone to the destination binary vector pMDC32 (*Pro35S::BdNRT2A*), pMDC43 (*Pro35S::GFP-BdNRT2A*) and pMDC83 (*Pro35S::BdNRT2A-GFP*) were performed. The vectors containing the different constructs were sequenced before transformation of *A. tumefaciens*. The *atnrt2.1-1* mutants (Ws background) (Filleur *et al.,* 2001) was transformed with each construct by the *in planta* method using the surfactant Silwet L-77 (Clough and Bent, 1998) and transformants were selected on 20 mg L^-1^ of hygromycin B (Sigma). Three independent homozygous mono-insertional T3 lines of were selected per construct, and over-expression was confirmed q-PCR. Roots of seven-days-old plantlets grown *in vitro* on Arabidopsis media (Duchefa Biochemie B.V; The Netherlands) were observed with the confocal microscope SP5 (Leica) or tested for the *BdNRT2A* overexpression by qRT-PCR.

### RFP fusions with *BdNRT3.2*

The cDNA of *BdNRT3.2* was amplified from genomic DNA using *EcoR1-BdNRT3_qL* and *BdNRT3_20_qR-Sal1*, and was cloned into the pGEMT-easy vector. Then, pGEMT containing BdNRT3.2 was digested by EcoR1 and Sal1, and the product of digestion was cloned by using T4 ligase, into pSAT6-RFP-C1 (NovoPro) and used for transitory expression of *p35S::RFP-BdNRT3.2* into mesophyll protoplasts of Arabidopsis.

### Directed mutagenesis of BdNRT2A

Directed mutagenesis of the conserved Ser specific to monocots (supplementary data S3) were performed using the protocol of ‘QuickChange II XL Site-Directed Mutagenesis’ (Stratagen). Sequence of *BdNRT2A* was amplified from pDONR207 containing *BdNRT2A* with the forward and reverse primers *BdNRT2A^S461D^*for BdNRT2A^S461D^ and the forward and reverse primers *BdNRT2A^S461A^*for BdNRT2A^S461A^. The amplification products were digested with *Dpn* I and then cloned into *E. coli* One Shot TOP10. The different constructs were sequenced, and those with desired mutations were re-introduced into pMDC43 (*Pro35S::GFP-BdNRT2A^S461D^*) and pMDC43 (*Pro35S::GFP-BdNRT2A^S461A^*) for transitory expression in Arabidopsis mesophyll protoplasts.

### Transfection of Arabidopsis protoplasts

The protoplast were isolated from seedlings of Arabidopsis and transfected as described in Zhai *et al.,* (2009). We used wild type genotype (Ws), *atnrt2.1-1, atnrt2-1.1xatnrt3.1,* and *atnrt2.1-1* overexpressing *Pro35S::GFPBdNRT2A* and *Pro35S::BdNRT2A-GFP* for isolation and transfection of protoplasts. Transfection were performed with pMDC43 containing *Pro35S::GFP-BdNRT2A* or pMDC83 containing *Pro35S::BdNRT2A-GFP*, and co-transfection were performed with pSAT6 containing the construction *Pro35S::GFPBdNRT3.2* and pMDC43 containing *Pro35S::GFP-BdNRT2A* or pMDC83 containing *Pro35S::BdNRT2A-GFP*. Co-transfection were also performed with pSAT6 (Citovsky *et al.,* 2006) containing the construction *Pro35S::GFPBdNRT3.2* and pMDC43 (*Pro35S::GFP-BdNRT2A^S461D^*) or pMDC43 (*Pro35S::GFP-BdNRT2A^S461A^*).

### Confocal imaging microscopy analyses

Confocal imaging microscopy analyses were performed using the Leica SP2 and SP5 microscope equipped with an argon laser (488 nm for GFP excitation and 543 nm for RFP). Emission was collected at 495-525 nm (GFP) and 580-650 nm (RFP). Autofluorescence was detected using the argon laser (488 nm) and emission was collected at 675-750nm. Images were processed in ImageJ.

### Root ^15^NO_3_^-^ influx

Root influxes of ^15^NO_3_^-^ were performed two weeks after growth in hydroponic conditions in order to measure HATS and ‘LATS plus HATS’ activities. First, plants were transferred to 0.1 mM CaSO4 for 1 min, then to a complete nutrient solution containing 0.2 mM of ^15^NO_3_^-^ for the HATS and 6 mM ^15^NO_3_^-^ for the ‘LATS plus HATS’ (atom% ^15^N : 99%) for 5 min and finally to 0.1 mM CaSO_4_ for 1 min. Roots were separated from the shoots immediately after the final transfer and frozen in liquid nitrogen. After grinding, an aliquot of the powder was dried overnight at 80°C and analyzed using a FLASH 2000 Elemental Analyzer coupled to an IRMS Delta IV (Thermo Fisher Scientific, Villebon, France). Influxes of ^15^NO_3_^-^ were calculated from the ^15^N content of the roots.

### NO_3_^-^ content

The NO_3_^-^ content was measured by a spectrophotometric method adapted from Miranda *et al.,* (2001) and described in David *et al.,* (2019). The principle of this method is a reduction of NO_3_^-^ by vanadium (III) combined with detection by the acidic Griess reaction.

### RNA extraction and qRT-PCR

Total RNAs were isolated using Trizol® reagent (Ambion, Life Technologies) and RT-qPCR were performed as described in David *et al.,* (2019).

### Statistical analyses

Statistical analyses were performed using one-way ANOVA and the means were classified using Tukey HSD test. (P<0.05)

## Results

### *BdNRT2A, BdNRT2B* and *BdNRT3.2* were induced in response to NO_3_^-^

We previously observed that the HATS activity in Brachypodium (Bd21-3 accession) decreased with increasing availability of NO_3_^-^ from 0.1 to 10 mM or 1 mM NH_4_NO_3_ supply, in correlation with the expression of *BdNRT2A/B* and *BdNRT3.2* in roots (David *et al*., 2019). The main component of the HATS activity in Arabidopsis, *AtNRT2.1,* is also repressed by high NO_3_^-^ and induced upon initial NO_3_^-^ supply (Filleur and Daniel-Vedele 1999; Lejay *et al*., 1999), similarly to *TaNRT2.1* which is induced by NO_3_^-^ in wheat (Yin *et al*., 2007). Since we found seven *BdNRT2* genes (Girin *et al.,* 2014), we further investigated the effect of 1 mM NO_3_^-^ re-supply (‘+N’) after 4 days of N deprivation (‘-N’) on root expression levels of the 5 members of the *BdNRT2* family which are most phylogenetically related to *AtNRT2.1*. *BdNRT2A* was induced by a factor 5 (Fig 1A), *BdNRT2B* and *BdNRT2F* were induced by a factor 2 (Fig 1B, Fig S1B), while *BdNRT2D* was extremely weakly expressed in both ‘-N’ and ‘+N’ conditions (Fig S1A) and *BdNRT2C* expression level was not modified by NO_3_^-^ availability (David *et al*., 2016). Besides, BdNRT2.E which is more phylogenetically related to AtNRT2.5 (Girin *et al.,* 2014) was repressed by NO_3_^-^ treatment (Fig S1C), while *BdNRT2.G*, phylogenetically related to *AtNRT2.7* (Girin *et al.,* 2014), was not expressed in roots (data not shown). Expression of *BdNRT3.2* was also induced by a factor 1.4 (Fig. 1C), whereas expression of *BdNRT3.1* was not affected by NO_3_^-^ availability (David *et al.,* 2016). Thus, *BdNRT2A* was the *BdNRT2* most induced in response to NO_3_^-^ induction (concomitantly with *BdNT3.2)*, an expression profile that fitted exactly with the characteristics of HATS activity.

**Figure 1.**
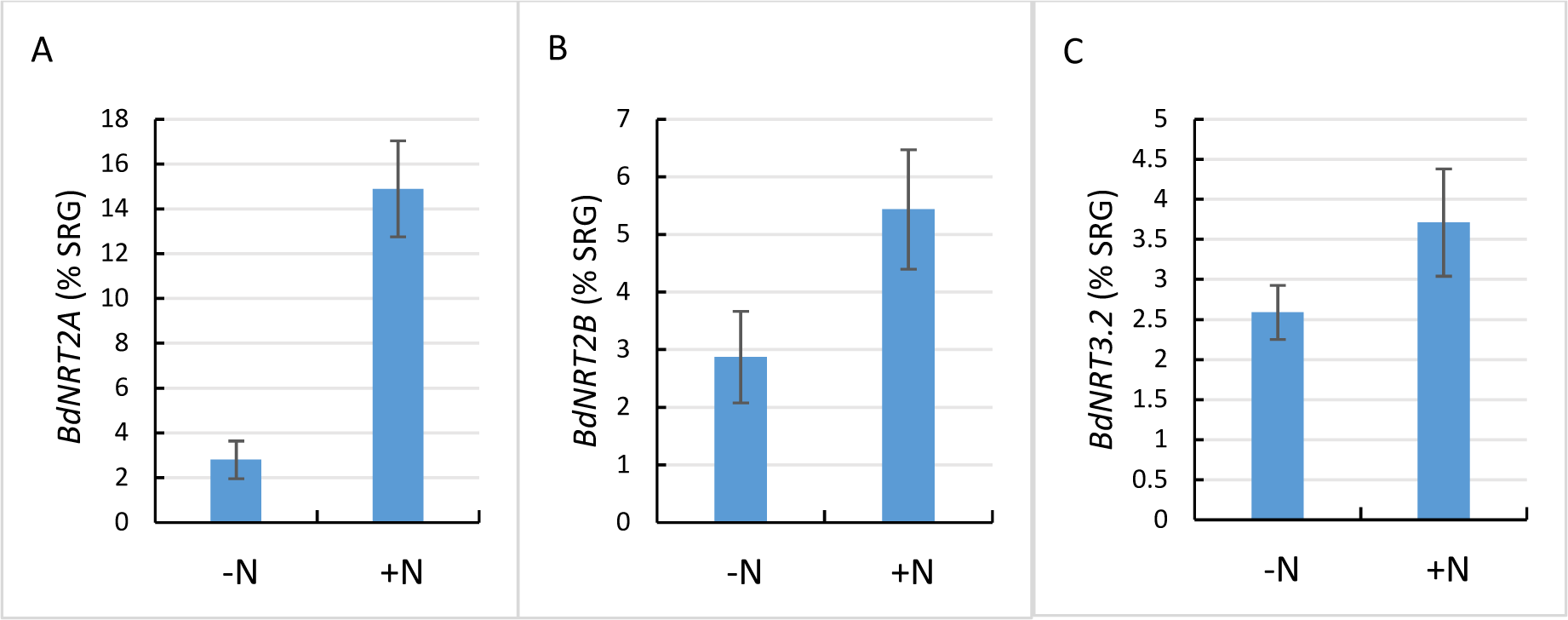
Root expression levels of *BdNRT2A* (1A), *BdNRT2B* (1B) and *BdNRT3.2* (1C) in response to 3h NO_3_^-^ re-supply (+N) after 4 days of N deprivation (-N). Gene expressions were quantified by qRT-PCR and were normalized to the level of a synthetic reference gene (SRG). *BdEF1α, BdUBC18* and *BdSAMDC* were used to compose the SRG. Values are means ± SD of 4 biological replicates.

### Co-expression of *BdNRT2A* and *BdNRT3.2* was required for an effective NO_3_^-^ transport in the heterologous expression system Xenopus oocytes

The two component system NRT2/NRT3 has been described as a hetero-oligomer in many species (Yong *et al*., 2010). Thus, we further investigated whether BdNRT2A needs to interact with BdNRT3.2 to transport NO_3_^-^. As already described in Orsel *et al*., (2006), we used the heterologous expression system in *Xenopus laevis* oocytes to express *BdNRT2A*, *BdNRT3.2* or both genes and then we measured the NO_3_^-^ uptake into oocytes. Only oocytes co-injected with cRNAs corresponding to BdNRT2A and BdNRT3.2 were able to accumulate ^15^NO_3_^-^ after 16 h of incubation in 0.5 mM Na^15^NO3, indicating that BdNRT2A and BdNRT3.2 interaction was required for an active transport system (Fig. 2) similarly to other species (Orsel *et al*., 2006).

**Figure 2.**
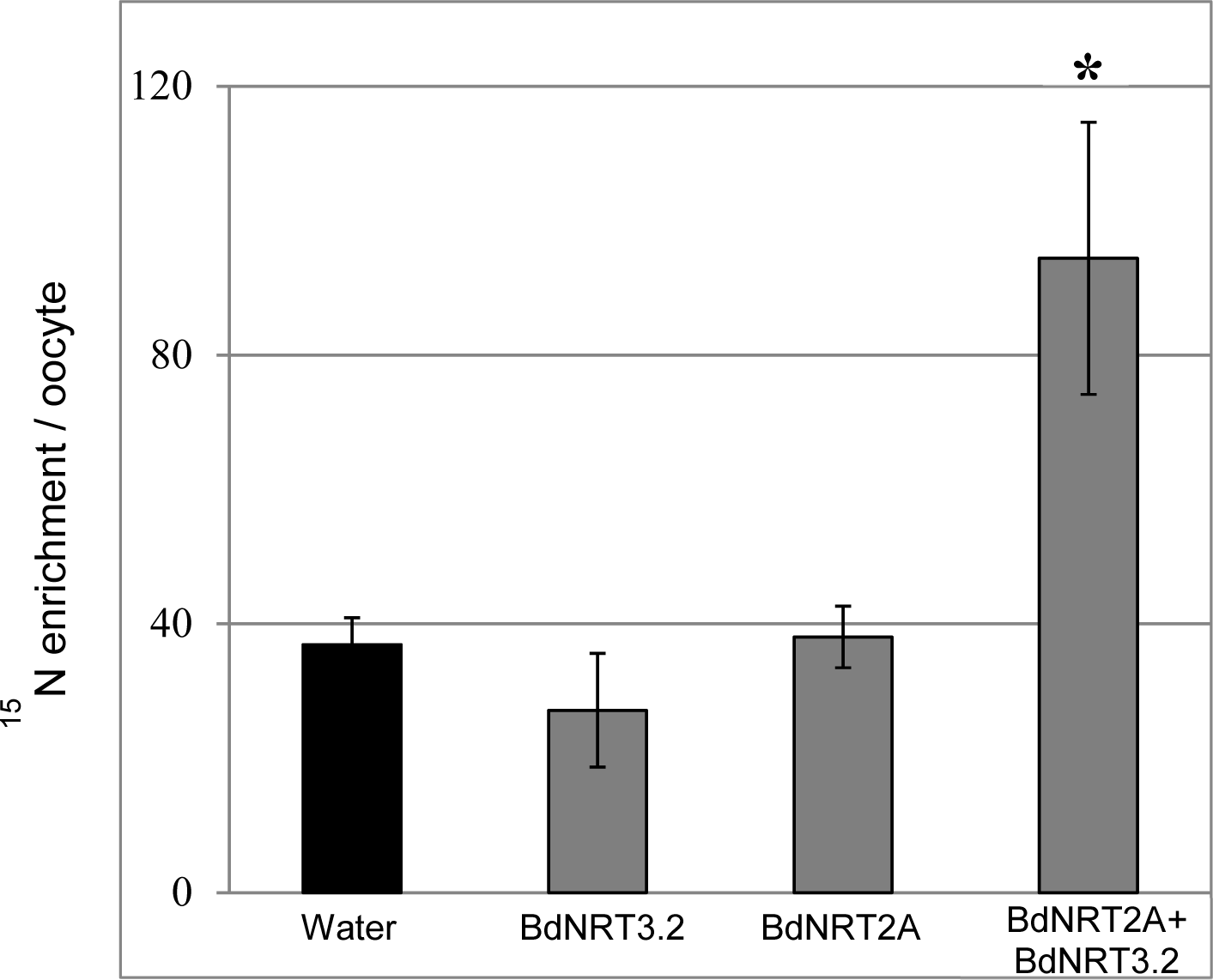
^15^NO_3_^-^ influx in heterologous expression system *Xenopus laevis* oocytes. *Xenopus laevis* oocytes were injected with nuclease-free water, *BdNRT2A, BdNRT3.2* or co-injected with *BdNRT2A* and *BdNRT3.2* cRNAs as indicated. After three days, oocytes were incubated in a solution enriched with 0.5 mM Na^15^NO_3_ (atom% ^15^N: 98%) and the ^15^N enrichment of individual oocytes was measured after 16h. The values are means ± SD of five replicates from a representative experiment. Asterisk indicates a statistically significant difference (Mann and Whitney, p<0.05).

### HATS activity and growth were lowered in Brachypodium mutants deficient in *BdNRT2A*

AtNRT2.1 is the main actor of HATS activity in Arabidopsis and a*tnrt2.1* mutants have a HATS activity reduced by 56% (Yin *et al*., 2007). In the double mutant a*tnrt2.1 atnrt2.2* more than 70% of the HATS was impaired (Filleur *et al*., 2001) while LATS activity was unaffected in both single and double mutants. In order to confirm that *BdNRT2A* and/or *BdNRT2B* are functional orthologs of *AtNRT2.1* and *AtNRT2.2* we searched for Brachypodium mutants affected in *BdNRT2A* and/or *BdNRT2B*. In the JGI Brachypodium collection no true T-DNA insertion mutant in *BdNRT2A* is available following our own testing experiments (Supplementary data S1). We used the NaN_3_ mutant collection from Versailles (Dalmais *et al*., 2013) to search for point mutations in the coding sequence of *BdNRT2A*. We identified seven independent lines corresponding to three silent and four non-silent point mutations leading to amino acid changes (Table 1). The SIFT scores for three of the non-silent point mutation were high (> 0.2) and thus with very low chance to impede the protein structure, but one mutation resulted in a stop codon (W248*) that likely resulted in a truncated protein deprived of 257 amino acids (out of 505 total amino acids) at the C-terminal. We further selected at the M_3_ generation one *bdnrt2a-W248*\* homozygous plant (*9.2*) on the one hand, and on the other hand two lines having lost by segregation the stop codon but potentially keeping other point mutations “azygotes” (az1 and az2).

**Table 1:** List of mutation sites identified in *BdNRT2A* by screening the Versailles NaN_3_ mutant collection (Dalmais *et al*. 2013). ^a^ Point mutation positions of in mutants relative to the starting ATG on the coding sequence. ^b^ Position of amino acid substitution in mutants are relative to the starting methionine of the encoded protein. ^c^ numbers are predictive score from the SIFT software (http://sift.bii.a-star.edu.sg/).

| Nucleic acid mutation <sup>a</sup> | Amino acid substitution <sup>b</sup> | SIFT<sup>c</sup> | Name of the mutant line chosen in this study |
| --- | --- | --- | --- |
| *G315A* | G105G | - | - |
| *T678C* | D226D | - | - |
| *C672T* | L224L | - | - |
| *G736A* | V336I | 0.2 |  |
| ***G744A*** | **W248\***<br>**STOP** | **0** | **9.2** |
| *C818T* | T273I | 0.46 | - |
| *G862A* | D288N | 0.9 | - |

We also obtained Brachypodium mutant lines using artificial micro RNA technology (amiRNA). The amiRNAs sequences were designed to target specifically *BdNRT2A* (*amiR n3*) or five out of the seven *BdNRT2 (2A, 2B, 2C, 2D, 2F)* (*amiR j2*). Expression of the amiRNA transgenes were verified in both independent transformed lines. *BdNRT2A* expression level was measured by qRT-PCR in order to verify the silencing effect of the amiRNA constructs. *BdNRT2A* and *BdNRT2B* mRNA level were decreased respectively by 26% and 66% in *amiRn3* line and by 45% and 10% in *amiRj2* line (Fig. S2).

The amiRNA mutant lines *amiRj2, amiRn3* and the TILLING mutant *9.2* line were grown in hydroponics under 0.2 mM NO_3_^-^ for three-week in order to study the impact of the decrease in *BdNRT2A* and *BdNRT2B* expression and of the presence of a truncated BdNRT2A protein on growth and NO_3_^-^ influx in comparison to WT and two azygotic (az) lines (az1, az2). All the mutants showed significant decreases in shoot/root ratio (Fig. 3A) resulting from a stronger decrease in shoot than in root biomass (Fig. S3). The *9.2* mutant line with a truncated BdNRT2A was the most affected with a decrease of 57% in shoot/root ratio, while decreases of 24% and 27% were observed for the shoot/root ratio in *amiRj2* and *amiRn3* respectively. Other developmental features such as increase in root length and decrease in tiller numbers were observed only in the mutant *9.2.* The mutant *9.2* showed a 1.7-fold increase in root length (Fig. 3B) and a 68% decrease in tiller number. The azygotes lines were not statistically different from the WT for the shoot/root ratio, root length and tiller numbers (Fig. 3A, 3B, 3C). However, the azygotes lines displayed reduced root and shoot biomasses compared to the WT, that could likely be due to the remaining point mutations in these lines, potentially disturbing the plant growth.

**Figure 3.**
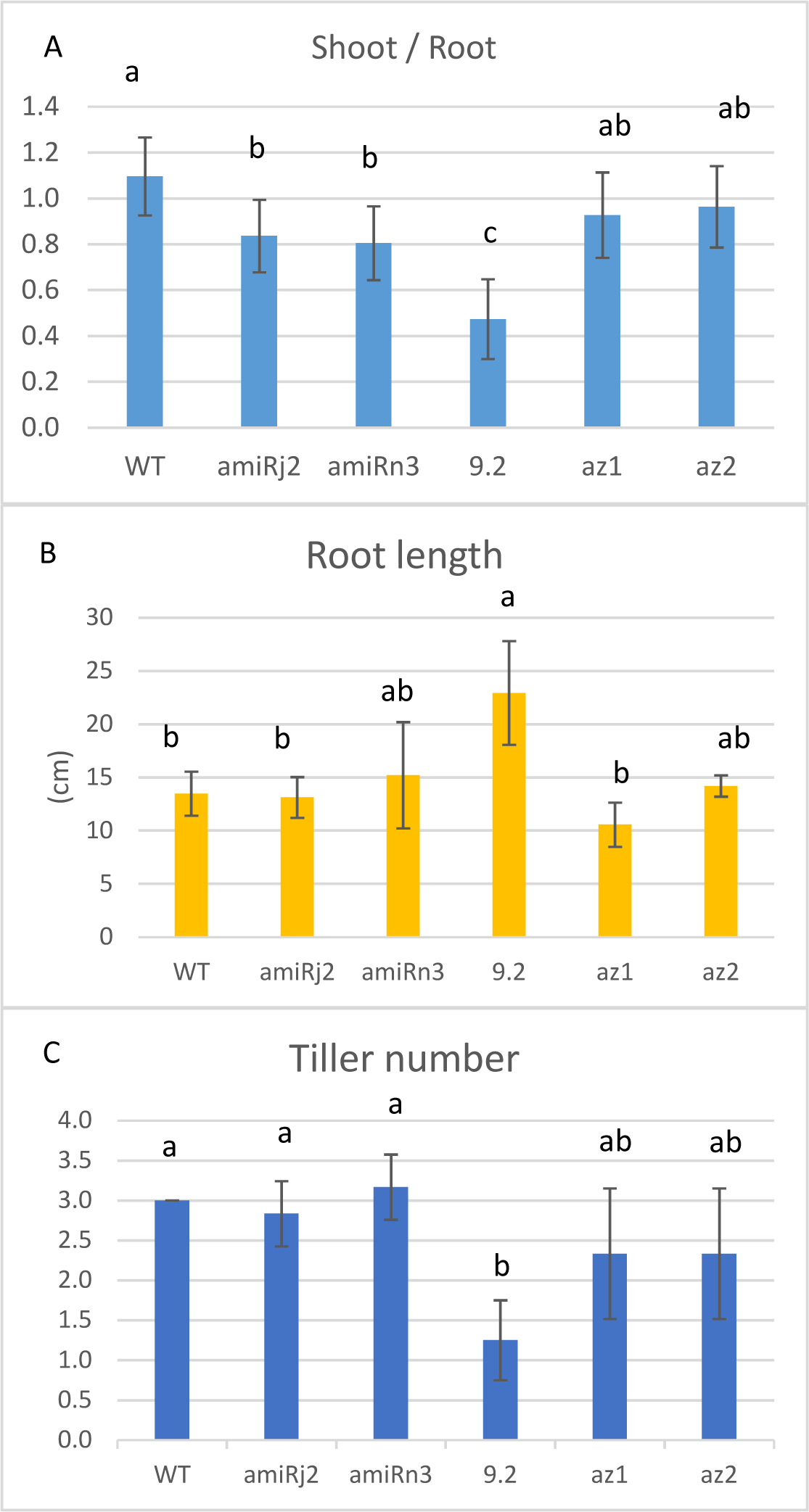
Growth phenotype of *bdnrt2a* mutants. Wild type and *bdnrt2a* mutants [two amiRNA lines : *amiRj2* and *amiRn3*, and one TILLING *bdnrt2a-W248*\* mutant (*9.2)*, and two azygotes lines az1 and az2] were grown in hydroponics with 0.2 mM NO_3_^-^ in controlled conditions, and the shoot/root biomass ratio (A), root length (B) and tiller number (C) were measured for 3 week-old plants. Values are means ± SD of 12 (A) and 6 replicates (B, C). Statistical analyses were performed using one-ANOVA and the means were classified using Tukey HSD test. (P<0.05).

HATS and the combined LATS and HATS activities were measured at 0.2 mM ^15^NO_3_^-^ and 6 mM ^15^NO_3_^-^ respectively on three-week-old plants grown in hydroponics. HATS was reduced significantly (up to 43% compared to WT) in the mutant *9.2*, and only a tendential decrease was observed in the *amiRj2* and *amiRn3* (up to 14% and 17% respectively compared to WT) (Fig. 4A) while the nitrate uptake at 6mM nitrate corresponding to the combined LATS and HATS was not significantly affected in all the mutants (Fig. 4B). The ^15^NO_3_^-^ influx of the azygotes lines was similar to the wild type for the ^15^NO_3_^-^ influx, suggesting that the decrease in ^15^NO_3_^-^ influx observed in the *9.2* mutant line was specifically due to the truncated BdNRT2A.

**Figure 4.**
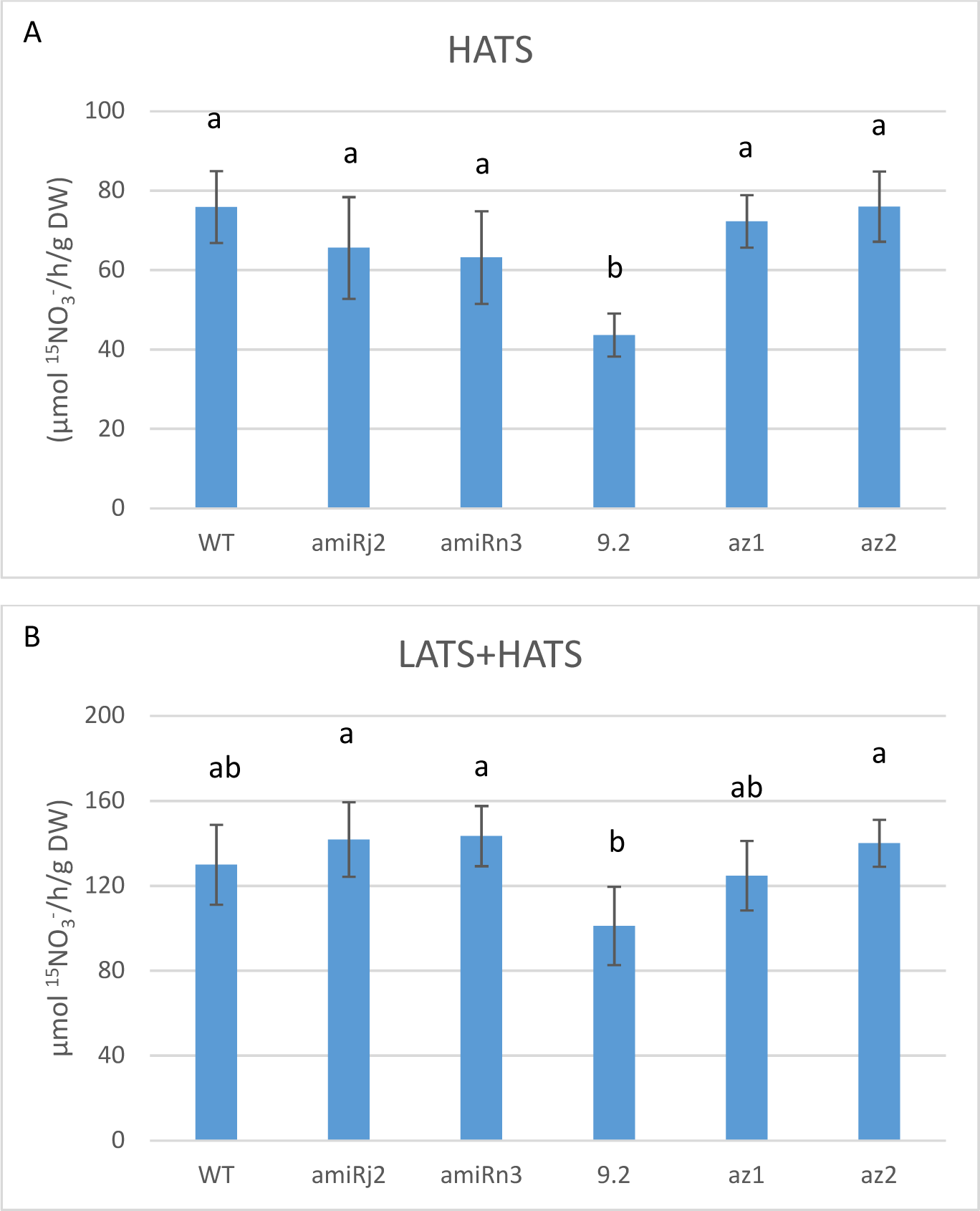
HATS and LATS activities in *bdnrt2a* mutants. Wild type and *bdnrt2a* mutants were grown in hydroponics with 0.2 mM NO_3_^-^ in controlled conditions, and HATS (A) and LATS (B) were measured at 0.2 mM ^15^NO_3_^-^ and 6 mM ^15^NO_3_^-^ respectively for 3 week-old plants. Values are means ± SD of 12 replicates. Statistical analyses were performed using one-way ANOVA and the means were classified using Tukey HSD test. (P<0.05).

Root total N contents were slightly but significantly decreased (by 6%) in the *9.2* mutant line in comparison to WT and az lines, while no differences have been observed for the amiR lines (Fig. 5A). In roots, nitrate content (Fig. 5B), total C content, C/N ratio were not changed for neither line compared to WT and az lines (S4A, S4B). Shoot nitrate content were not significantly affected either (Fig S4C). All together, these results showed that the *9.2* mutant line was the most affected line for HATS activity and growth, likely due to the loss of function of a truncated BdNRT2A. These results confirmed the conserved functional role for BdNRT2A similar to AtNRT2.1, since *atnrt2.1-1* mutant shows a reduced HATS activity when grown on 0.2 mM of nitrate as sole nitrogen source, resulting in a lower biomass compared to the wild type (Filleur *et al*., 2001).

**Figure 5.**
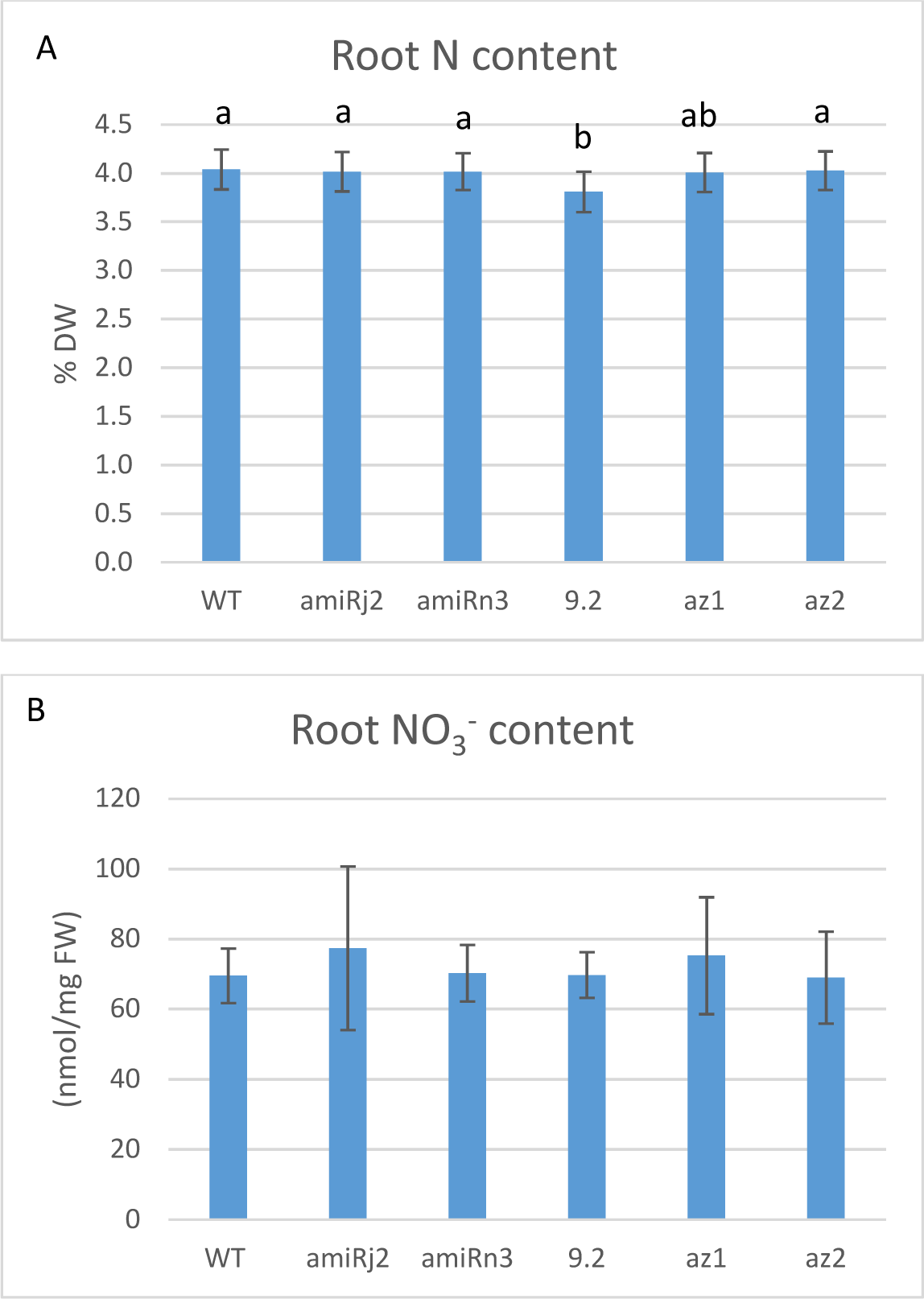
Total N and nitrate contents in roots of *bdnrt2a* mutants. Wild type and *bdnrt2a* mutants were grown in hydroponics with 0.2 mM NO_3_^-^ in controlled conditions, and the total N content (A) and root nitrate content (B) were measured for 3 week-old plants. Values are means ± SD of 12 replicates. Statistical analyses were performed using one-way ANOVA and the means were classified using Tukey HSD test. (P<0.05).

We further performed hydroponic cultures using this mutant in order to study the role of BdNRT2A under a range of NO_3_^-^ supplies from 0.02 mM to 10 mM nitrate for 3 weeks. The *9.2* mutant line showed a significant increase in root/shoot ratio whatever the nitrate supply, and an increase in root length under 0.2 and 10mM nitrate, while the tiller number was reduced only under 10 mM nitrate (Fig. S5A, S5B, S5C). These results suggest that BdNRT2A modulated Brachypodium growth even under a large range nitrate supply.

### Overexpression of *BdNRT2A* in *atnrt2.1-1* was not sufficient for a functional complementation

In order to further investigate the function of BdNRT2A *in planta*, we produced *atnrt2.1-1* plants overexpressing *BdNRT2A* translationally fused or not with Green Fluorescent Protein (GFP) coding sequence (in C or N terminal position) under the control of *35S* promoter. For each construct, three independent over-expressing lines were selected (Fig. S6). Then, functional complementation test was performed on plants grown under hydroponic conditions at 0.2 mM of nitrate. Shoot biomass of the complemented lines were reduced as compared to the WT line, and similar to the *atnrt2.1-1* mutant (Fig. 6A). Root ^15^NO_3_^-^ influx was measured at 0.2 mM NO_3_^-^ and showed that HATS activity was not restored to the wild type level in any of the tested complemented lines (Fig. 6B). Moreover, a cytosolic sub-cellular localization of BdNRT2A fused to GFP indicated that BdNRT2A was not targeted at the pm in the *atnrt2.1-1* lines overexpressing *Pro35S::BdNRT2A-GFP* or *Pro35S::GFP-BdNRT2A*, for both C and N terminal GFP fusions (Fig. 6C). Unexpectedly, these results demonstrate that heterologous overexpression of *BdNRT2A* in *atnrt2.1-1* was not sufficient for a functional complementation of the mutant, likely due to the lack of BdNRT2A targeting to the pm.

**Figure 6.**
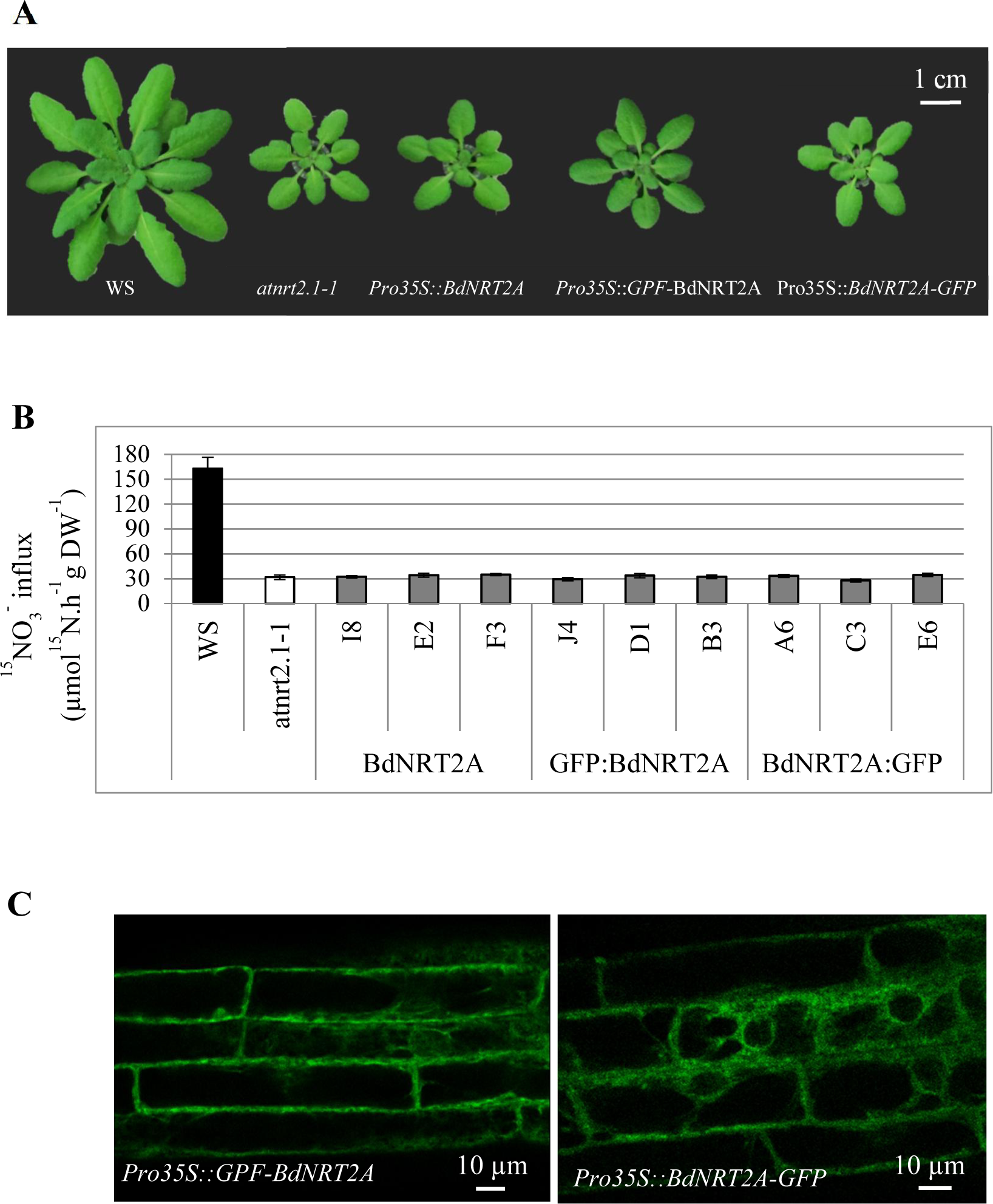
Overexpression of *BdNRT2A* in *atnrt2.1-1* mutant. A) Rosette area of wild type (WS), *atnrt2.1-1* and three lines expressing *Pro35S::BdNRT2A*, *Pro35S::GFPBdNRT2A* or *Pro35S::BdNRT2A-GFP* in the *atnrt2.1-1* mutant. Plants were grown hydroponically for 40 days on a nutrient solution containing 0.2 mM NO_3_^-^. (B) ^15^NO_3_^-^ influx (HATS), measured at the external concentration of 0.2 mM ^15^NO_3_^-^. Values are means of 9 replicates ± SE. (C) Sub-cellular localization of BdNRT2A fused to GFP in leaves of *atnrt2.1-1* mutant lines expressing *Pro35S::GFP-BdNRT2A*. For each construct, a representative picture of the 3 independent transgenic lines is presented for each construct.

### BdNRT2A was targeted to plasma membrane in presence of BdNRT3.2 but not in the presence of AtNRT3.1

In Arabidopsis, HATS is mediated by two component systems. The interaction between NRT2 and NRT3 proteins is required for the pm targeting of the complex and for an active NO_3_^-^ transport (Wirth *et al*., 2007). However, we did not observe BdNRT2A targeting to the pm in *atnrt2.1-1* complemented with *BdNRT2A*, suggesting that BdNRT2A could not interact with AtNRT3.1. Besides, we observed that co-expression of *BdNRT2A* and *BdNRT3.2* was required for an effective NO_3_^-^ transport in the heterologous expression system *Xenopus* oocytes, validating the two-component system in Brachypodium. Then, to further investigate the sub-cellular localization of the BdNRT2A/BdNRT3.2 complex in Arabidopsis by using another expression system, we used transient expression in mesophyll protoplasts from Arabidopsis. We transfected mesophyll protoplasts from Arabidopsis seedlings with *Pro35S::GFP-BdNRT2A* and/or *Pro35S::RFP-BdNRT3.2*.

Interestingly, when *Pro35S::GFP-BdNRT2A* was expressed transiently in protoplasts from the single mutant *atnrt2.1*, the localization of BdNRT2A fusion protein was cytosolic (Fig. 7A), similarly to what was observed in leaves or in protoplasts obtained from an *atnrt2.1-2* mutant stably transformed with *Pro35S::GFP-BdNRT2A* line (Fig. 6C, Fig.S7). Conversely, when *Pro35S::GFP-BdNRT2A* and *Pro35S::RFP-BdNRT3.2*) were co-expressed transiently in protoplasts of *atnrt2.1-1* mutant, BdNRT2A and BdNRT3.2 fusion proteins were co-localized at the pm (Fig. 7B). Moreover, when mesophyll protoplasts from *atnrt2.1-1 Pro35S::GFP-BdNRT2A* line (B3) were transfected with *Pro35S::RFP-BdNRT3.2,* we observed a pm colocalization of BdNRT2A and BdNRT3.2 fusion proteins (Fig. 8), corroborating that BdNRT2A could not interact with AtNRT3.1 and that the interaction with BdNRT3.2 allows its targeting to the pm. Besides, when we used transient expression of *Pro35S::GFP-BdNRT2A* and *Pro35S::BdNRT2A*-*GFP* in *Nicotiana benthamiana* leaves, we also observed a cytosolic localization for BdNRT2A fusion protein in epidermal cells (Fig. S8). These results revealed a species-specific interaction between NRT2 and NRT3 proteins that ensures BdNRT2A targeting to the pm.

**Figure 7.**
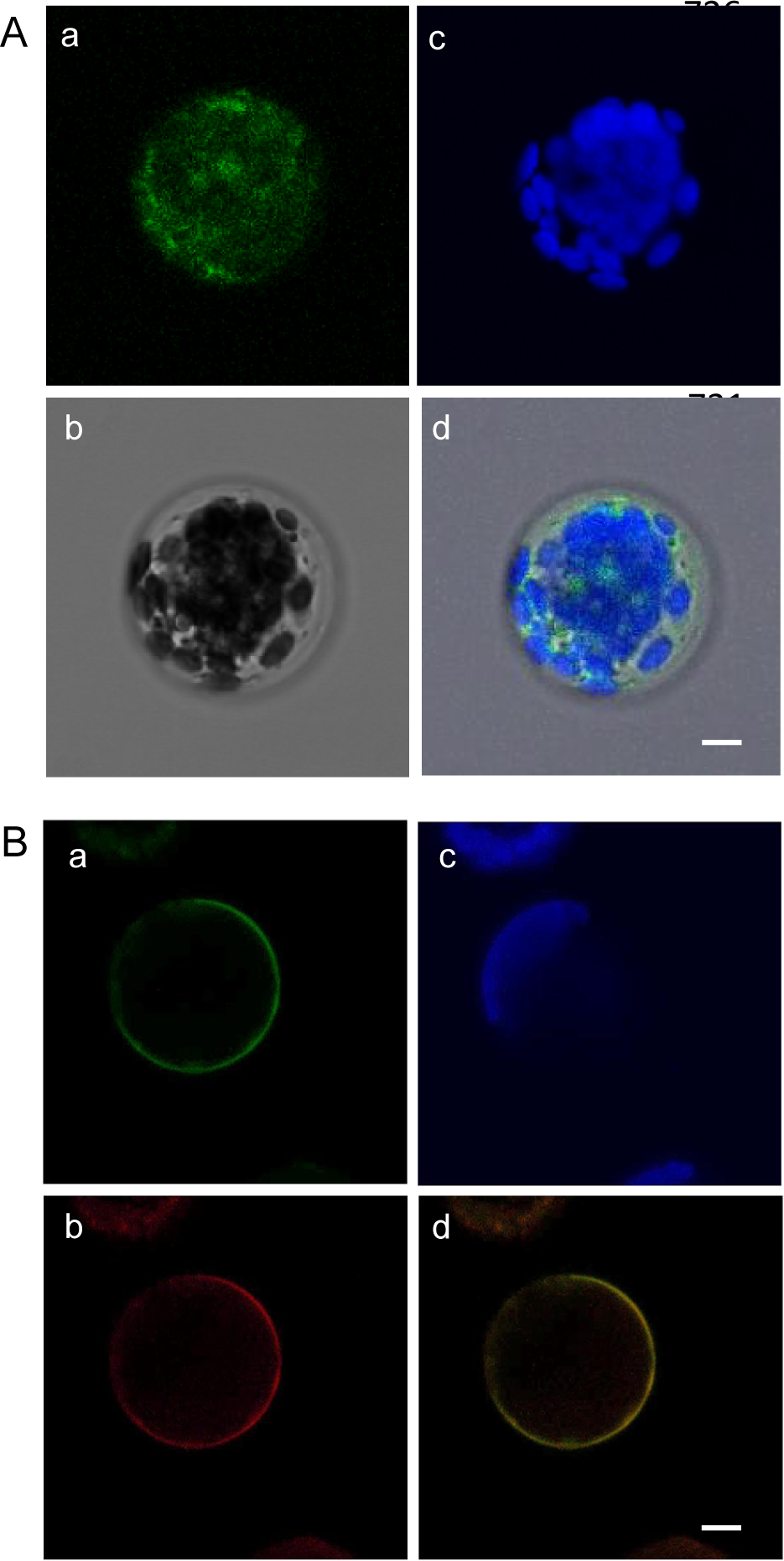
Subcellular localization of BdNRT2A-GFP and BdNRT3.2-RFP fusion proteins in Arabidopsis mesophyll protoplasts. Confocal images from *atnrt2.1-2* protoplasts transiently expressing *Pro35S::GFP-BdNRT2A* (A), or transiently co-expressing *Pro35S::GFP-BdNRT2A* and *Pro35S::RFP-BdNRT3.2 (*B). Different images are presented in Fig (A) : GFP fluorescence(a), bright-field image (b), chlorophyll auto-fluorescence indicating position of chloroplasts (c), and merged (d). Different images are presented Fig (B) : GFP fluorescence (a), RFP fluorescence (b), chlorophyll auto-fluorescence indicating position of chloroplasts (c) and merged (d). Scale bar = 5 µm.

**Figure 8.**
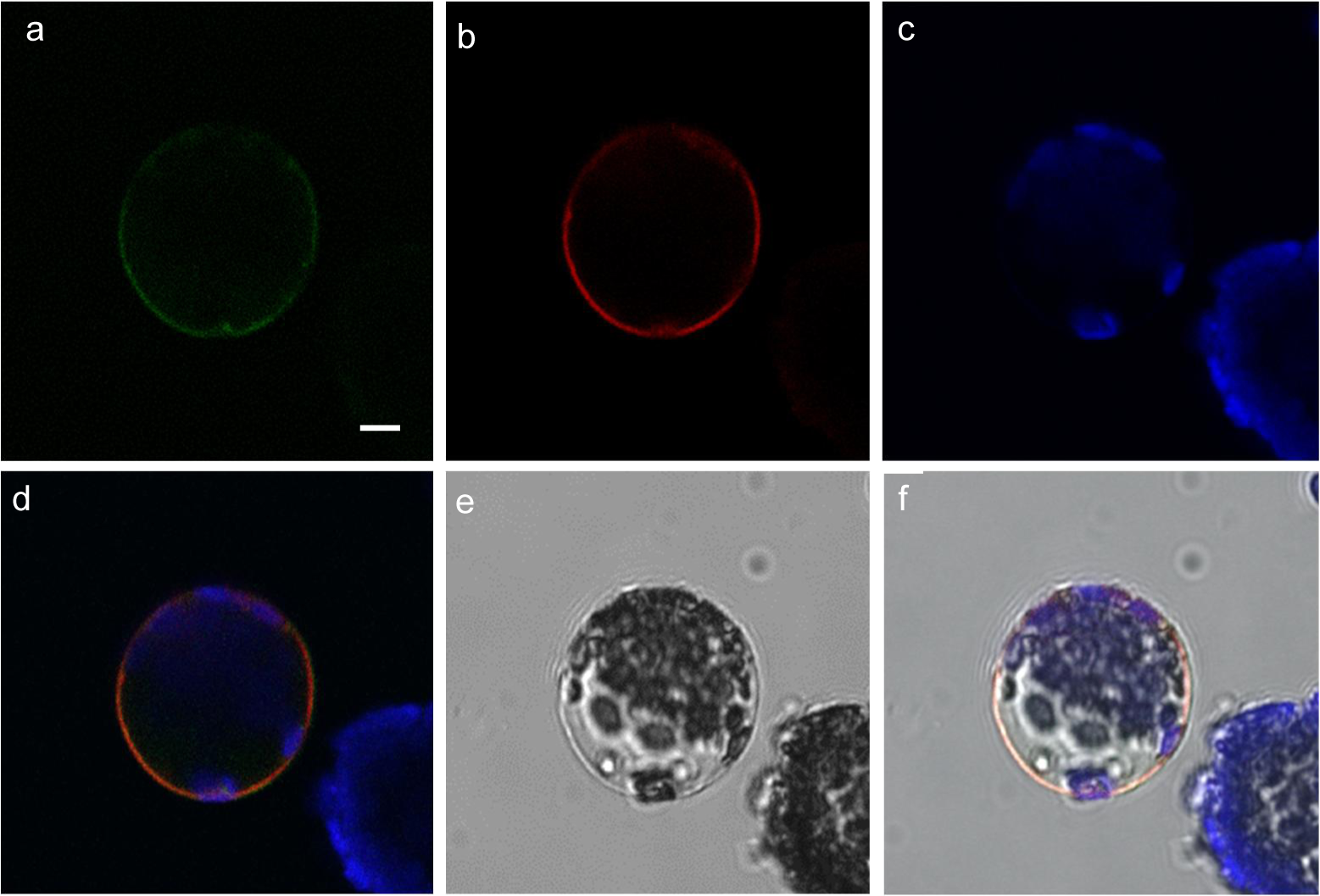
Subcellular localization of BdNRT2A-GFP and BdNRT3.2-RFP fusion proteins in Arabidopsis mesoplyll protoplasts. Confocal images of protoplasts from the *atnrt2.1-2* mutant line B3 expressing *Pro35S::GFP-BdNRT2A* and transiently expressing *Pro35S::RFP-BdNRT3.2*. Six images are presented : GFP fluorescence (a), RFP fluorescence (b), chlorophyll auto-fluorescence indicating position of chloroplasts (c), GFP and RFP merged with autofluorescence (d), bright field (e), and merged of all images (f). Scale bar = 5µm.

### The conserved BdNRT2A S461 residue is not required for the interaction between BdNRT2A and BdNRT3.2 in Brachypodium

The molecular mechanism involved in NRT2 and NRT3 interaction is not completely elucidated. However, in barley the interaction between HvNRT2.1 and HvNRT3.1 is impaired when HvNRT2.1 Ser463 in the C-terminus is changed to alanine that mimics a non-phosphorylated residue (Ishikawa *et al*., 2009). This residue is conserved in NRT2 proteins from monocotyledons and algae only (Jacquot *et al*., 2017) (supplementary data S3) and not in dicotyledons as for AtNRT2.1. We investigated whether this Ser residue is required for the interaction between BdNRT3.2 and BdNRT2A similarly to barley. We thus co-transfected mesophyll protoplasts of *atnrt2 atnrt3.1* with *Pro35S::RFP-BdNRT3.2* and *Pro35S::GFP-BdNRT2A^S461A^* or *p35S::GFP-BdNRT2A^S461D^*that mimicked a non phophorylated and a constitutively phosphorylated Ser, respectively. Surprisingly both mutagenized constructs allowed the targeting of BdNRT2A/BdNRT3.2-GFP/RFP fusion proteins at the pm, suggesting that the phosphorylated state of S461 (S461A or S461D) had no impact on the capacity of BdNRT2A to interact with BdNRT3.2 (Fig. 9) unlike in barley. Thus, the absence of the conserved S461 specific to monocots in AtNRT2.1 seems to not be responsible for the impaired pm targeting of BdNRT2A in Arabidopsis, that is likely due to lack of interaction between BdNRT2A and AtNRT3.1.

**Figure 9.**
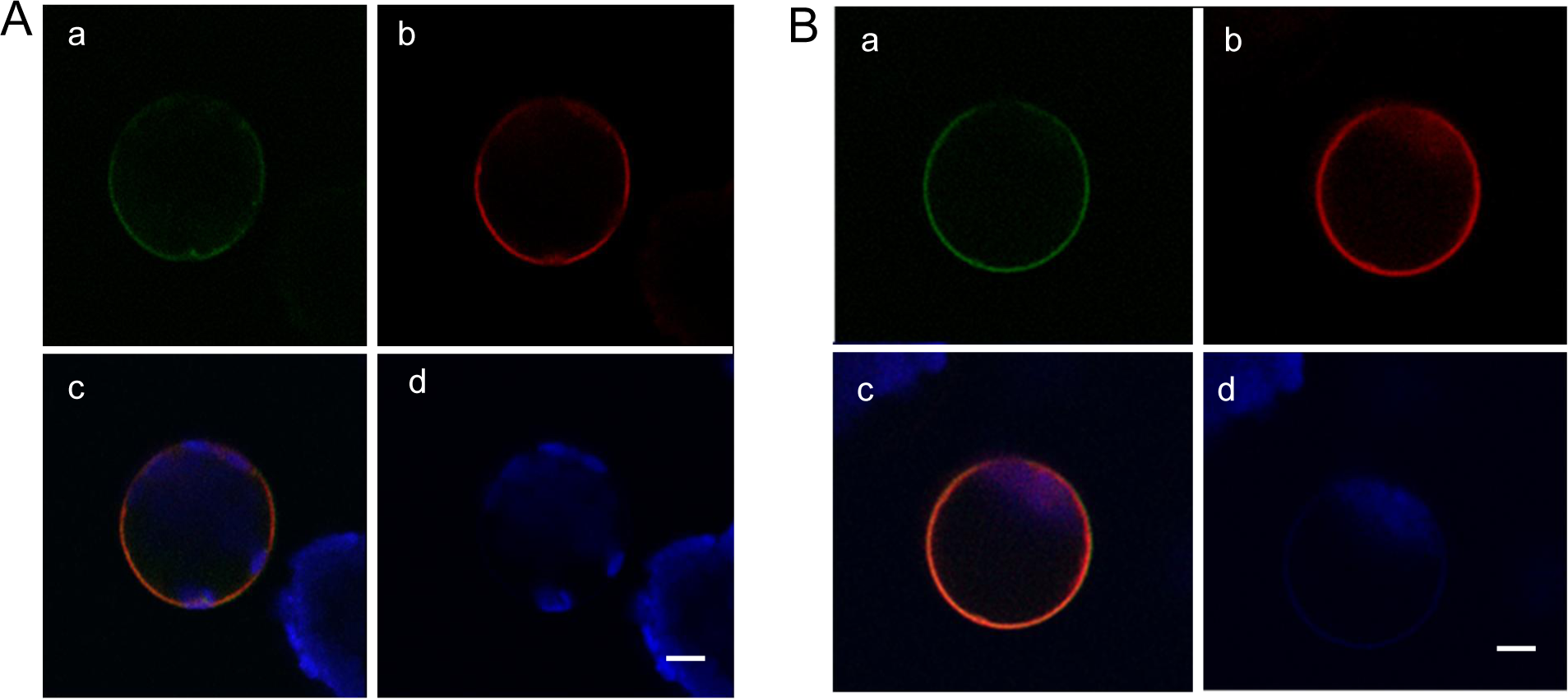
Subcellular localization of BdNRT2A^S461A^-GFP or BdNRT2A^461D^-GFP and Bd NRT3.2-RFP fusion proteins in Arabidopsis mesophyll protoplasts. Confocal images of protoplasts *of atnrt2.1-2xatnrt3.1* double mutant transiently co-expressing *Pro35S::GFP-BdNRT2A^S461A^* and *Pro35S::RFP-BdNRT3.2 (*A), or transiently co-expressing *Pro35S::GFP-BdNRT2A^S461D^*and *Pro35S::RFP-BdNRT3.2* (B). Four images are presented : GFP fluorescence (a), RFP fluorescence (b), merged (c), and chlorophyll auto-fluorescence indicating position of chloroplasts (d). Scale bar = 5 µm.

These results did not allow to explain the impaired targeting of BdNRT2A to pm in absence of BdNRT3.2 in Arabidopsis, but emphasized the complexity of this protein-protein interaction and need for further investigation of this interaction.

## Discussion

HATS and LATS activities for NO_3_^-^ in Brachypodium were already characterized in part in our previous paper (David *et al.,* 2019). HATS activity was regulated by N availability and correlated to the expression level of BdNRT2A/2B and BdNRT3.2 suggesting these genes are functional orthologous of AtNRT2.1/2.2 and AtNRT3.1, respectively, and are good candidates involved in HATS activity (David *et al*., 2019). In order to complete the characterization of the molecular basis of the HATS, we generated two *bdnrt2* mutants (*n3* and *j2* lines) using amiRNA strategy and made a TILLING analysis that allowed to select one NaN_3_ induced mutant (line *9.2*) with a truncated protein at the C terminus (amino acid W248*). The growth phenotype observed for *bdnrt2a*^W248^* (line *9.2*) was reminiscent to that of the insertional mutant *atnrt2.*1 in Arabidopsis (Orsel *et al.,* 2004), with reduced shoot/root ratio, shoot biomass and tiller number, and increased root length, likely due to lack of a functional protein BdNRT2A. Moreover, the NO_3_^-^ HATS activity was also reduced (up to 43%) in *bdnrt2a*^W248^* similar to the decrease in HATS observed in *atnrt2.1* (Orsel *et al.,* 2004, Li *et al.,* 2007). A less marked decrease in HATS was observed for *amiRj2* and *amiRn3* mutants (up to 14% and 17% respectively), in which slight decreases in *BdNRT2A* and *BdNRT2B* transcript levels were observed (but were not statistically significant compared to wild type) and then likely induced only slight decreased BdNRT2A and BdNRT2B protein levels. Besides, a slight reduction of shoot/root ratio and shoot biomass were observed for *amiRj2* and *amiRn3,* indicating that growth of these mutants was limited in correlation with the slight decrease in HATS. Interestingly, *amiRn3* seemed to be more reduced in *BdNRT2B* expression than in *BdNRT2A* but its phenotype was not different from *amiRj2*, suggesting that *BdNRT2A* is the main actor of HATS activity.

A slight but significant decrease in root N content was observed in *bdnrt2a*^W248^* (line *9.2*), but no change was observed for root and shoot NO_3_^-^ content in this mutant, while N and NO_3_^-^ content were not changed in *amiRj2* and *amiRn3*. On the contrary a 3 fold lower NO_3_^-^ content has been described in shoots of *atnrt2.1* accompanied by a slight decrease in root and shoot N content (Orsel *et al*., 2004). We previously observed that at low NO_3_^-^ supply (for NO_3_^-^ < 2 mM) and its assimilation products were used in priority for growth, and NO_3_^-^ storage in shoot occurred only under higher NO_3_^-^ supply in Brachypodium (David *et al*., 2019). This strategy seemed to be different from Arabidopsis, and that could explain why a lower NO_3_^-^ uptake in *bdnrt2a*^W248^* was accompanied by a reduced growth without changes in NO_3_^-^ content. We did not include in our study a *bdnrt2a* insertional mutant, although one line is displayed as available in the insertional mutant collection the JGI Brachypodium collection (studied in Wang *et al*., 2019), because we proved after thorough verifications that the T-DNA insertion was not placed into our gene of interest BdNRT2A (supplementary data S1).

Next, we overexpressed *BdNRT2A* with or without GFP-tag in the *atnrt2.1-1* mutant background to investigate if BdNRT2A can functionally complement *atnrt2.1-1*, but surprisingly the functional complementation was not observed, since the different complemented lines showed a reduced growth phenotype and a lower HATS activity, similar to *atnrt2.1-1*. This lack of functional complementation of *atnrt2.1-1* mutant by overexpression of *BdNRT2A* is probably due to the subcellular localization of the BdNRT2A-GFP fusion protein, that was cytosolic and not in the pm as initially expected. Our results remind the diffused fluorescence throughout the cell, that was observed when AtNRT2.1 fused to GFP was constitutively expressed in *atnar2.1-1* (also named *atnrt3.1*) background, while in wild type background, it was clearly associated with pm (Orsel *et al.,* 2007). The authors hypothesized that AtNRT3.1 is involved in the stability of AtNRT2.1 and possibly through the pm targeting process, leading to NO_3_^-^ uptake, as it was demonstrated later (Okamoto *et al.,* 2006, Wirth *et al*., 2007, Kotur *et al*., 2012). Besides, we confirm that BdNRT2A and BdNRT3.2 are responsible for the two-component system of HATS using a heterologous expression system in oocytes, similarly to other species. Consequently, we hypothesized that, in contrast to NpNRT2.1 (Filleur *et al*., 2001), BdNRT2A could not interact with AtNRT3.1 in the *atnrt2.1-1* line overexpressing *BdNRT2A* with or without GFP-tag and that was subsequently verified using transfection of Arabidopsis mesophyll protoplasts in either a wild type or an *atnrt2.1-1* background. When transfection was performed with *BdNRT2A* fused to *GFP, BdNRT2A* was cytosolic, and on the contrary, the co-expression of *BdNRT2A* and *BdNRT3.2,* fused to *GFP* and *RFP* respectively, allowed the targeting of the BdNRT2A/BdNRT3.2 to the pm in either an *atnrt2.1-1* or an *atnrt2.1-1 x atnrt3.1* background. Post-translational regulations of AtNRT2 occur in response to variations of N supply (Engelsberger and Schulze 2012; Menz *et al*., 2016), and phosphorylation of AtNRT2.1-S^28^ is crucial for the ATNRT2.1 stability in response to NO_3_^-^ limitation (Zou *et al.,* 2020). Recently, S501 in the C-terminus of AtNRT2.1 was found to be phosphorylated in NH_4_NO_3_ conditions leading to inactivation of AtNRT2.1 and decrease of NO_3_^-^ influx, but not to a dissociation of the AtNRT2.1/AtNRT3.1 complex (Jacquot *et al*., 2020). Interestingly this S501 residue is not conserved in monocotyledons, such as Brachypodium, and replaced by a Gly. On the other hand, another residue Ser is present in monocotyledonous and not conserved in dicotyledons, and this residue S463 was required for interaction of HvNRT2.1 and HvNRT3.1 (Ishikawa *et al.,* 2009). Thus, we hypothesized that this residue could play an important role in the species specificity we observed for NRT2.1/NRT3.1 interaction. Using the transfection of protoplasts, we observed no effect of the S461 substitution to S461A and S461D mimicking respectively constitutively non-phophorylated and phophorylated residues. The plasmalemic sub-cellular co-localization of BdNRT2A and BdNRT3.2 was not changed by the substitution of Ser461 suggesting that it was not involved in the regulation of BdNRT2A and BdNRT3.2 interaction in Brachypodium unlike in barley.

In conclusion, we demonstrated that BdNRT2A has a major role in NO_3_^-^ uptake at low N availability, and that BdNRT2A and BdNRT3.2 are the main component of the HATS in Brachypodium. The functional complementation experimentations in Arabidopsis suggested that a monocotyledons/dicotyledons species specificity exists for the NRT2A and NRT3 interaction, that was not explained by the presence of specific residue S461 conserved only in monocotyledons.

## Acknowledgements

We thank Hervé Ferry and Michel Burtin for taking care of the plants at the IJPB. We thank Rozenn Le-Hir for her help and advices in confocal manipulation and we thank the IJPB’s Plant Observatory technical plateforms for their support. We thank the TILLING plateform EpiTrans, INRA, 2018. EPIgenomics and TRANSlational Research Facility, doi: 10.15454/1.5572407597184844E12. We are gratefull to Christian Meyer for critical reading of the manuscript. This work was supported in part by Saclay Plant Sciences (ANR-10-LABX-0040), and by ANR NiCe (ANR-17-CE20-0021).

## Authors contribution

SFM, LCD designed and planned the experiments at IJPB. LCD, PB, SFM, MG, TG performed or participated to the various experiments. LCD performed the oocyte ^15^ N uptake experiments at the John Ines Center under the supervision of AJM. TG planned the production of the amiRNA and PB generated the transformed plants. LCD selected the tilling mutants under the supervision of MD and AB. SFM, PB, TG and LCD performed the ^15^N influx experiments. SFM, PB and LCD performed the protoplasts transfection, and confocal analyses. AM performed the ^15^N and N analyses. SFM, LCD, TG analyzed the data. TG, FDV and AK contributed to the design of this study. SFM wrote the manuscript and AK, TG, LCD, FDV, AJM, AM participated to its critical reading.

## List of primers

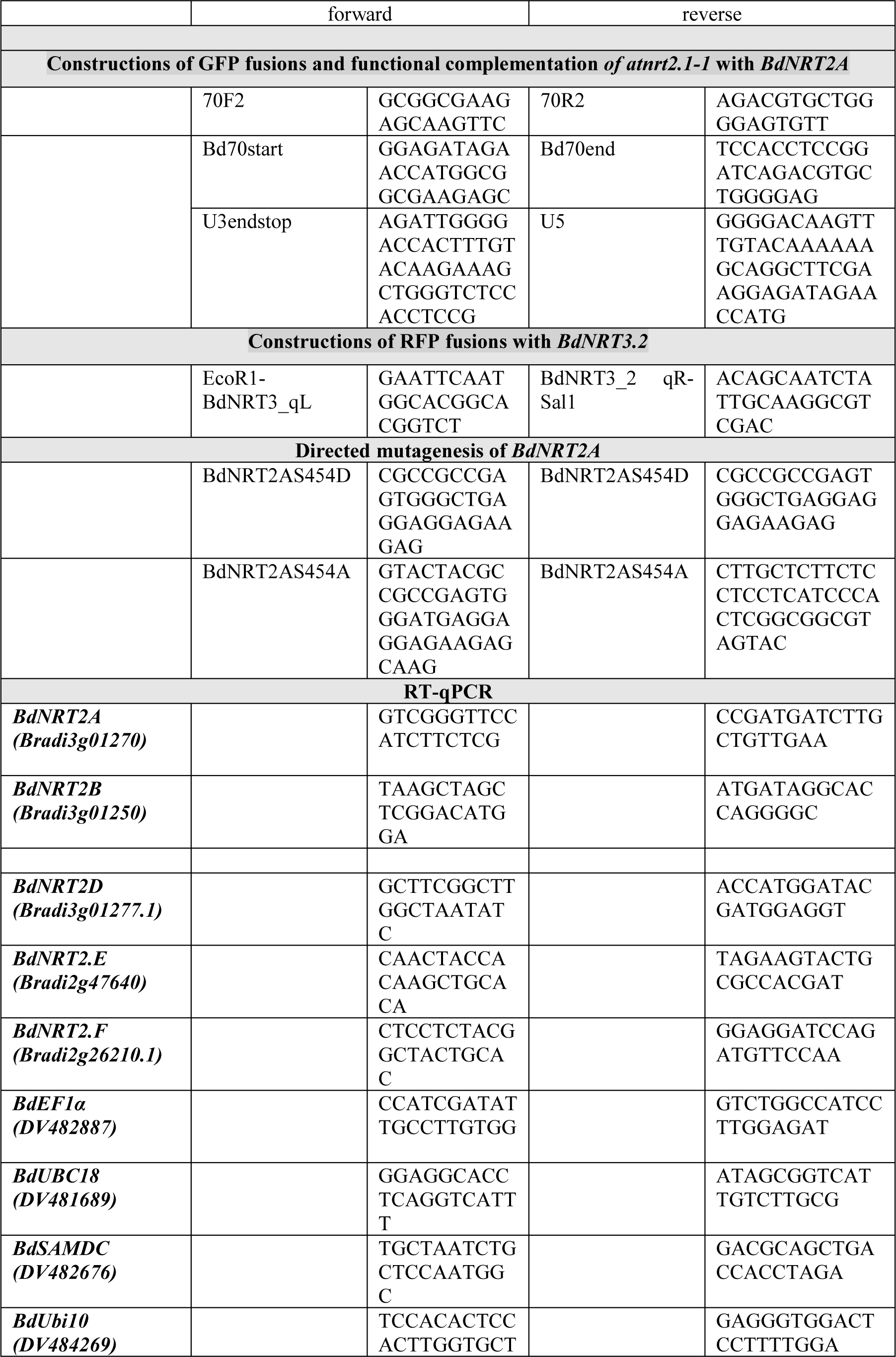
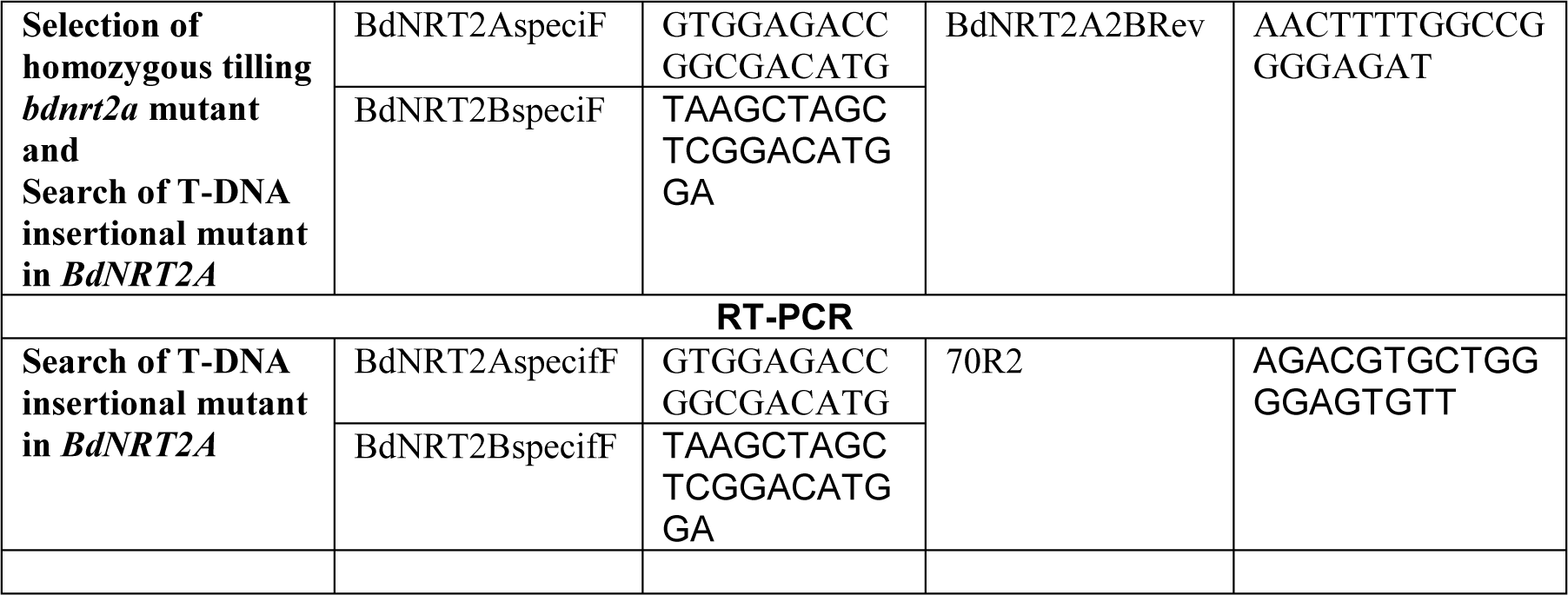

---

**Figure S1.**
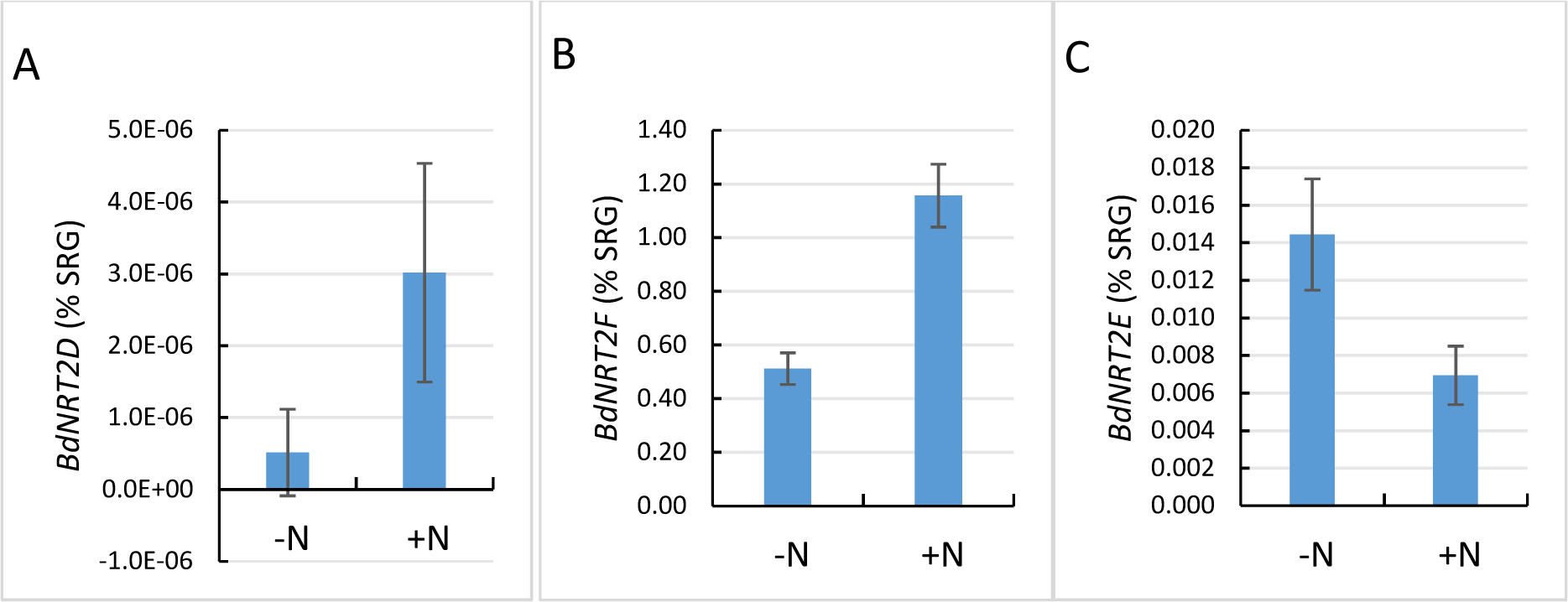
Root expression levels of *BdNRT2D* (S1A), *BdNRT2F* (S1B) and *BdNRT2.E* (S1C) in response to 2h NO_3_^-^ re-supply (+N) after 4 days of N deprivation (-N). Genes expressions were quantified by qRT-PCR and were normalized to the level of a synthetic reference gene (SRG). *BdEF1α, BdUBC18* and *BdSAMDC* were used to compose the SRG. Values are means ± SD of 4 biological replicates.

**Figure S2.**
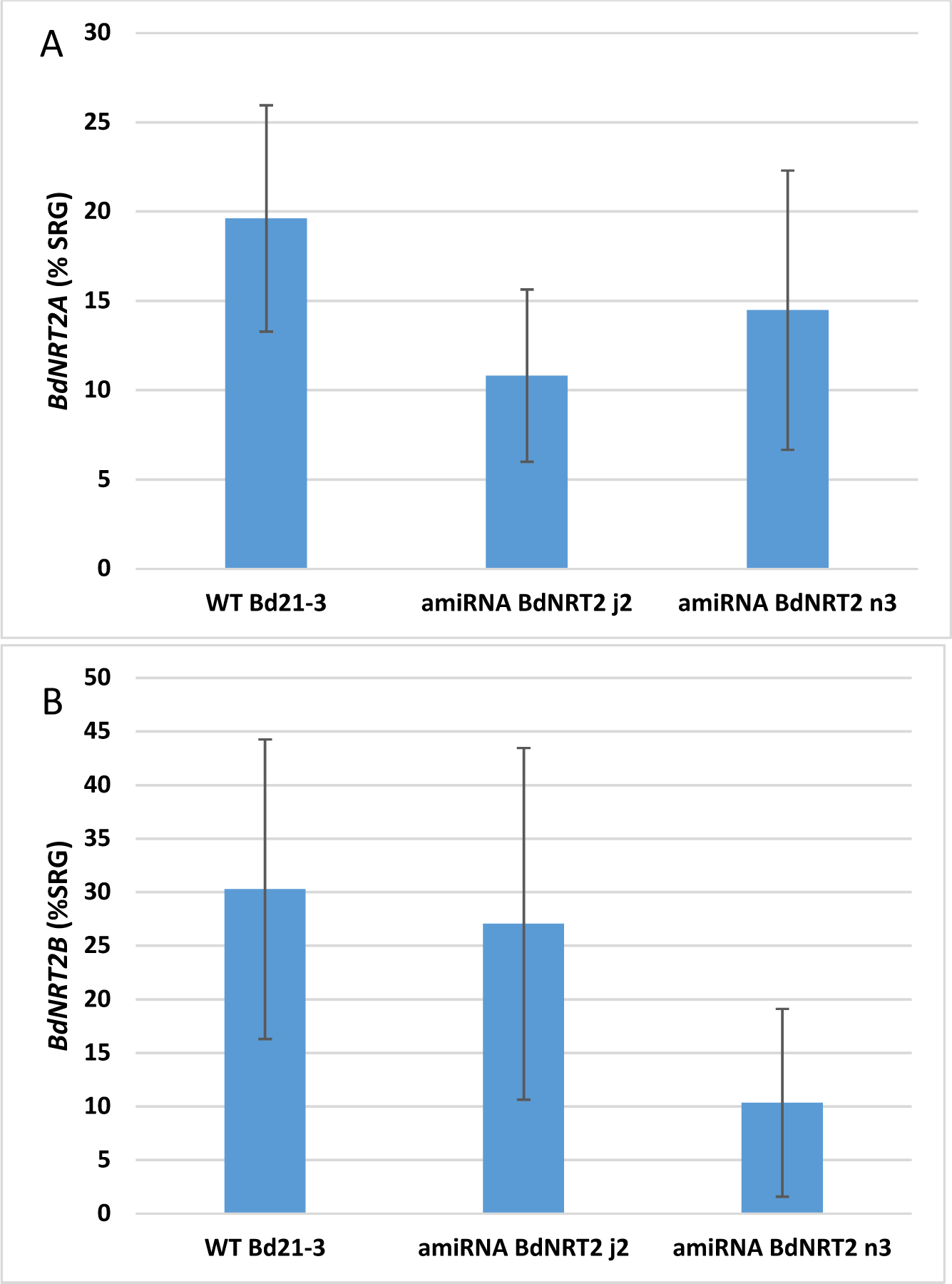
Root expression levels of *BdNRT2A* (4A) and *BdNRT2B* (4B) in amiRNA mutants *amiRj2, amiRn3.* Gene expressions were quantified by qRT-PCR and were normalized to the level of a synthetic reference gene (SRG). *BdUBI10* and *BdSAMDC* were used to compose the SRG. Values are means ± SD of 4 biological replicates.

**Figure S3.**
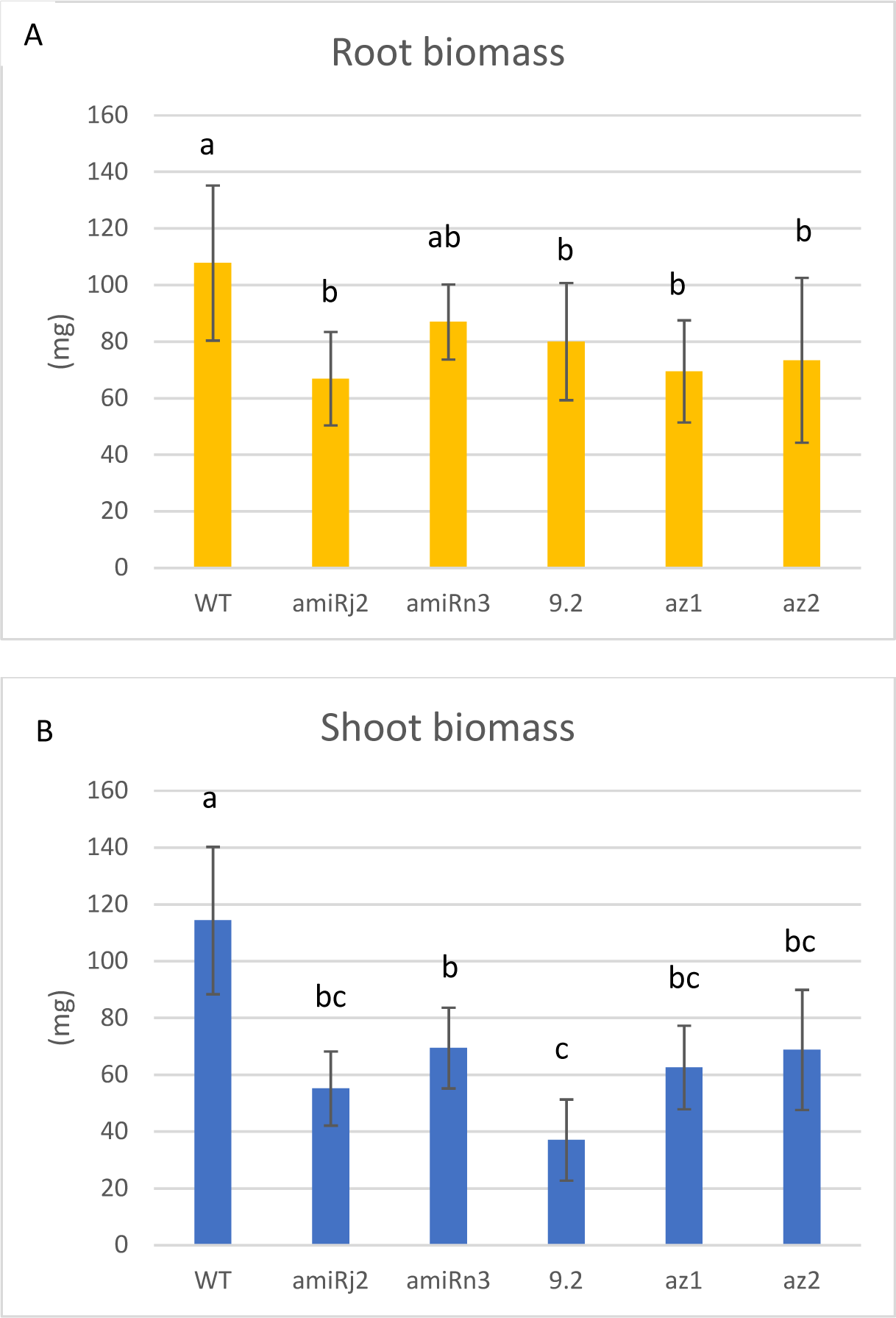
Growth phenotype of *bdnrt2a* mutants. Wild type and *bdnrt2a* mutants were grown in hydroponics with 0.2 mM NO_3_^-^ in controlled conditions, and the root biomass ratio (A) and shoot biomass (B) were measured for 3 week-old plants. Values are means ± SD of 12 replicates. Statistical analyses were performed using one-way ANOVA and the means were classified using Tukey HSD test. (P<0.05).

**Figure S4.**
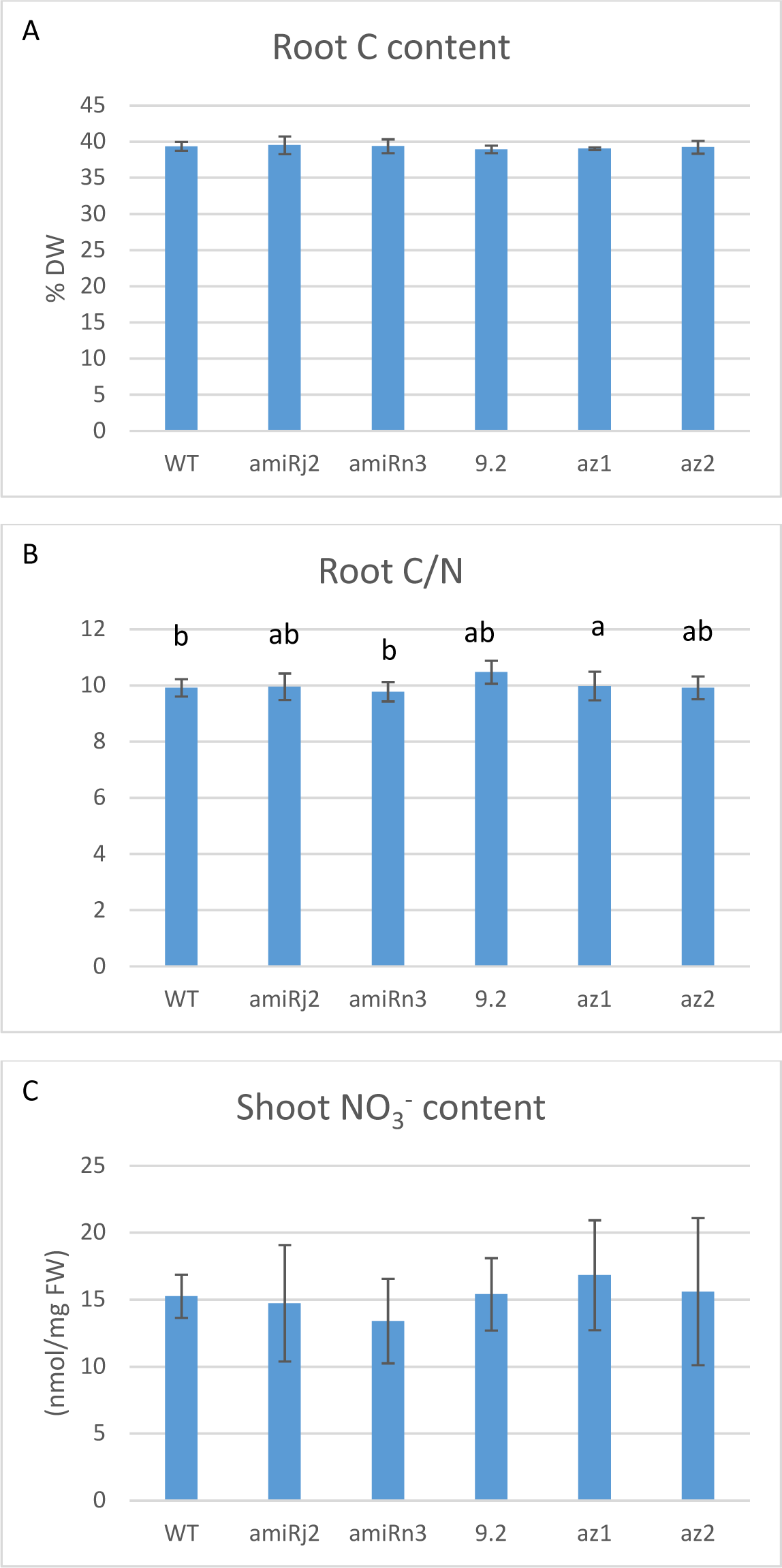
Total C and C/N in roots and shoot nitrate content in *bdnrt2a* mutants. Wild type and *bdnrt2a* mutants were grown in hydroponics with 0.2 mM NO_3_^-^ in controlled conditions. Total C content and C/N in roots (A, B) and shoot nitrate content (C) were measured for 3 week-old plants. Values are means ± SD of 12 replicates. Statistical analyses were performed using one-way ANOVA and the means were classified using Tukey HSD test. (P<0.05).

**Figure S5.**
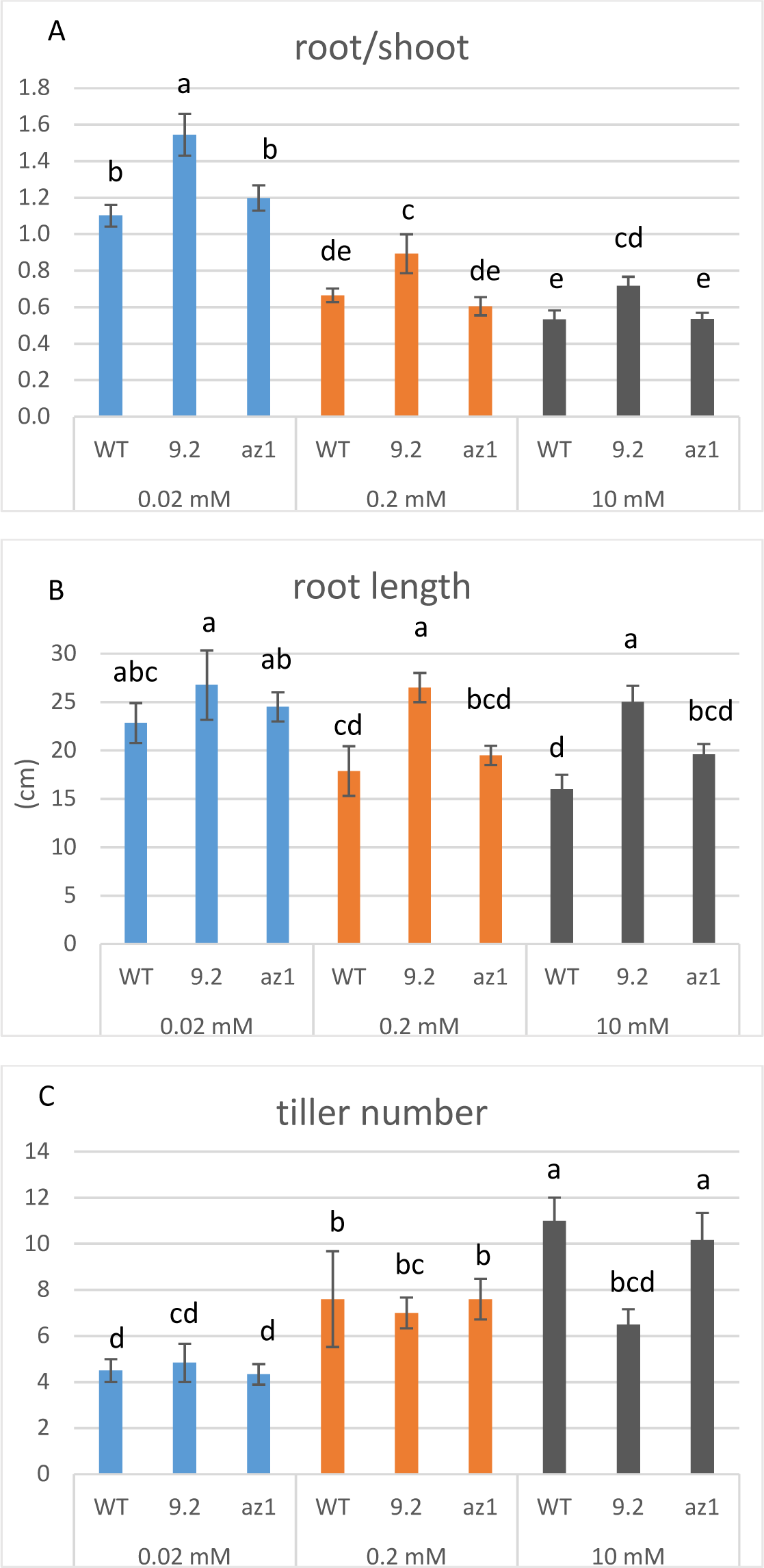
Growth phenotype of *bdnrt2a* mutants under varying nitrate supply. Wild type and *bdnrt2a* mutants were grown in hydroponics with 0.02, 0.2 and 10 mM NO_3_^-^ in controlled conditions. Shoot/root ratio (A), root length (B), tiller number (D) and were measured for 3 week-old plants. Values are means ± SD of 12 replicates. Statistical analyses were performed using one-way ANOVA and the means were classified using Tukey HSD test. (P<0.05).

**Figure S6.**
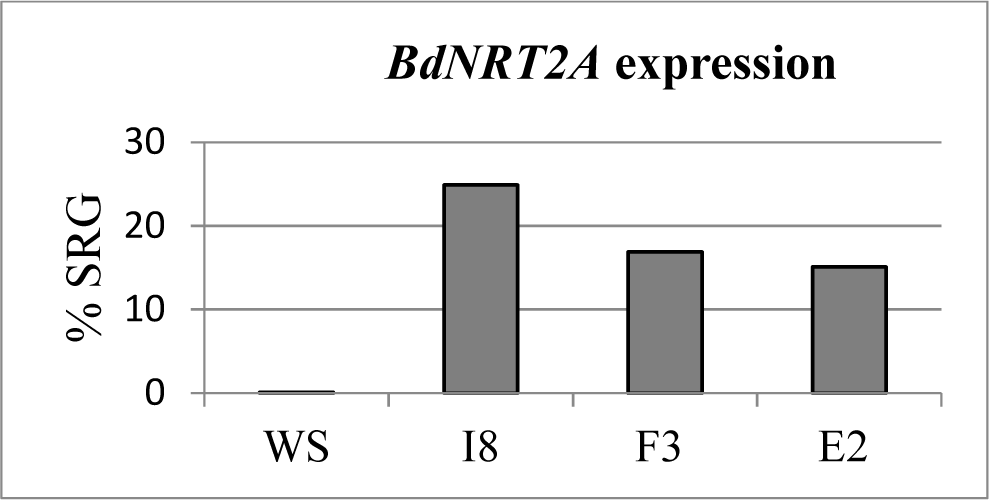
Overexpression of *BdNRT2A* in *atnrt2.1-1* mutant. Overexpression of *BdNRT2A* has been quantified in *atnrt2.1-1* lines overexpressing *Pro35S::BdNRT2A*. Gene expressions were quantified by qRT-PCR and was normalized to the level of a synthetic reference gene (SRG). *BdACT* and *BdPP2A3* were used to compose the SRG.

**Figure S7.**
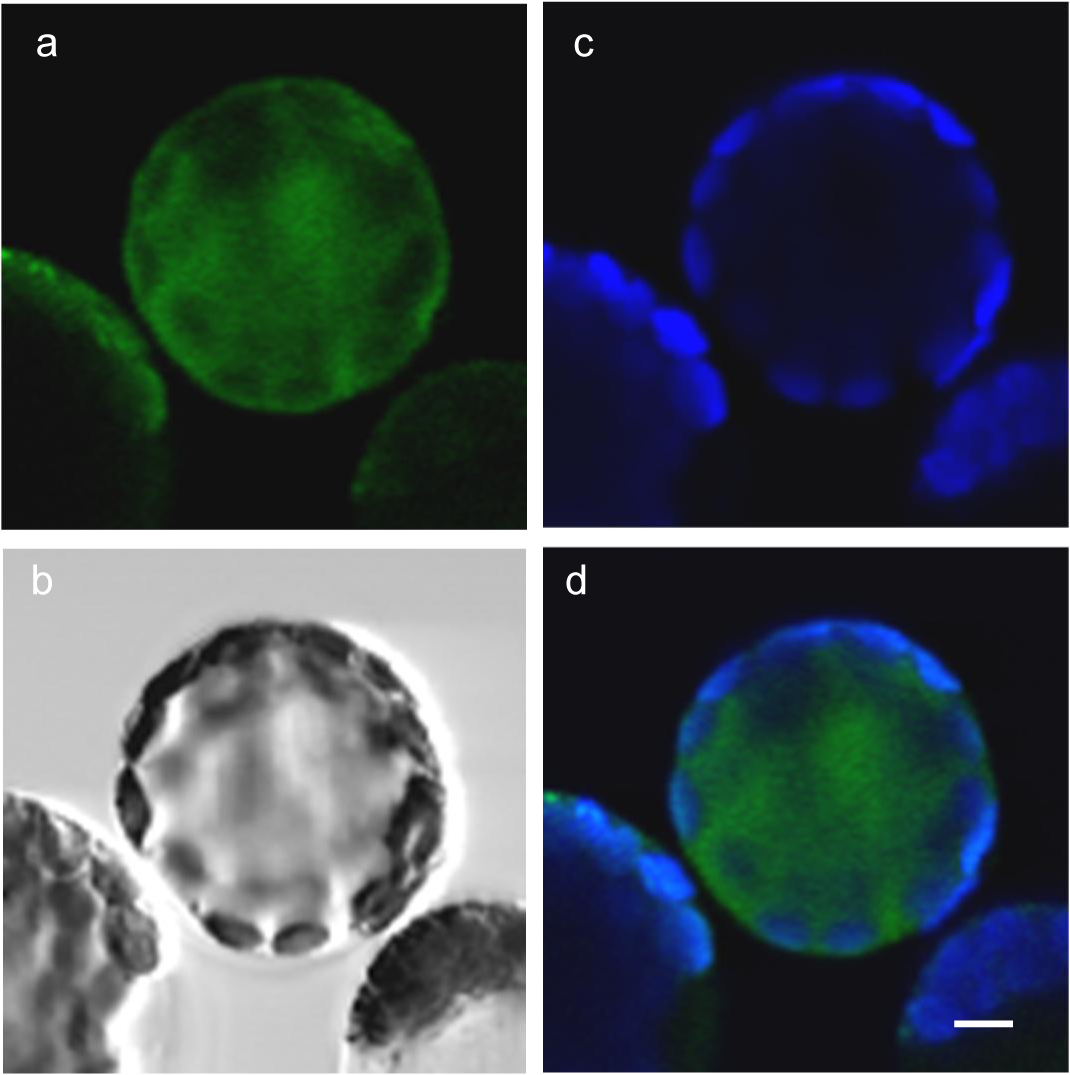
Subcellular localization of BdNRT2A-GFP fusion proteins in Arabidopsis mesophyll protoplasts in *atnrt2.1-2* mutant line B3. Confocal images from protoplasts of *atnrt2.1-2* mutant line B3 expressing *Pro35S::GFP-BdNRT2A*. Four images are presented : GFP fluorescence (a), bright field (b), chlorophyll auto-fluorescence indicating position of chloroplasts (c), and merged (d). Scale bar = 5 µm.

**Figure S8.**
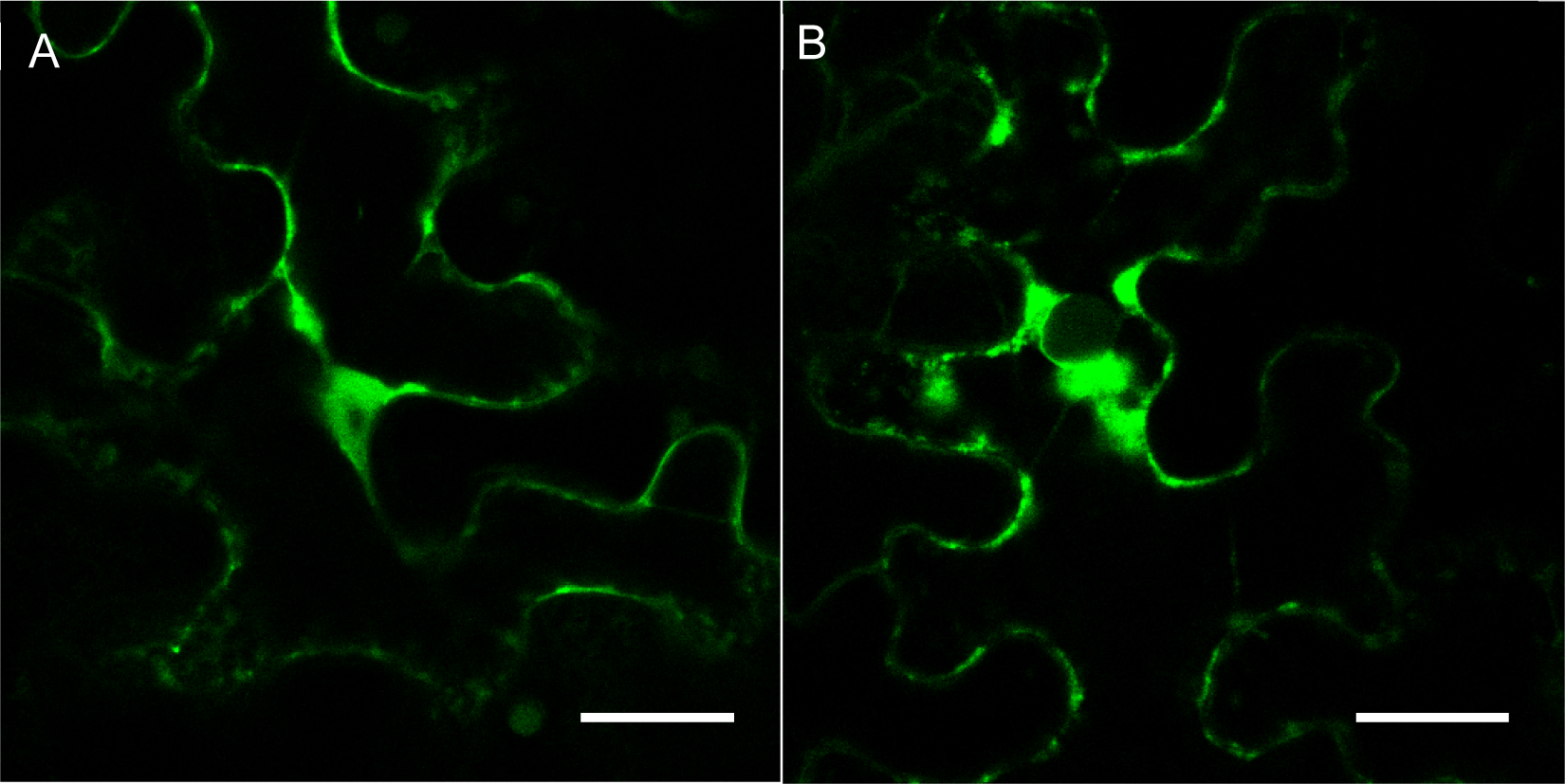
Subcellular localization of BdNRT2A-GFP fusion protein in *Nicotiana benthamiana* leaves. Confocal images from mesophyll tobacco cell transiently expressing *Pro35S::GFP-BdNRT2A* (A), *Pro35S::BdNRT2A-GFP (*B). GFP fluorescence are presented. Scale bar = 50 µm.

**Supplementary data S1:**
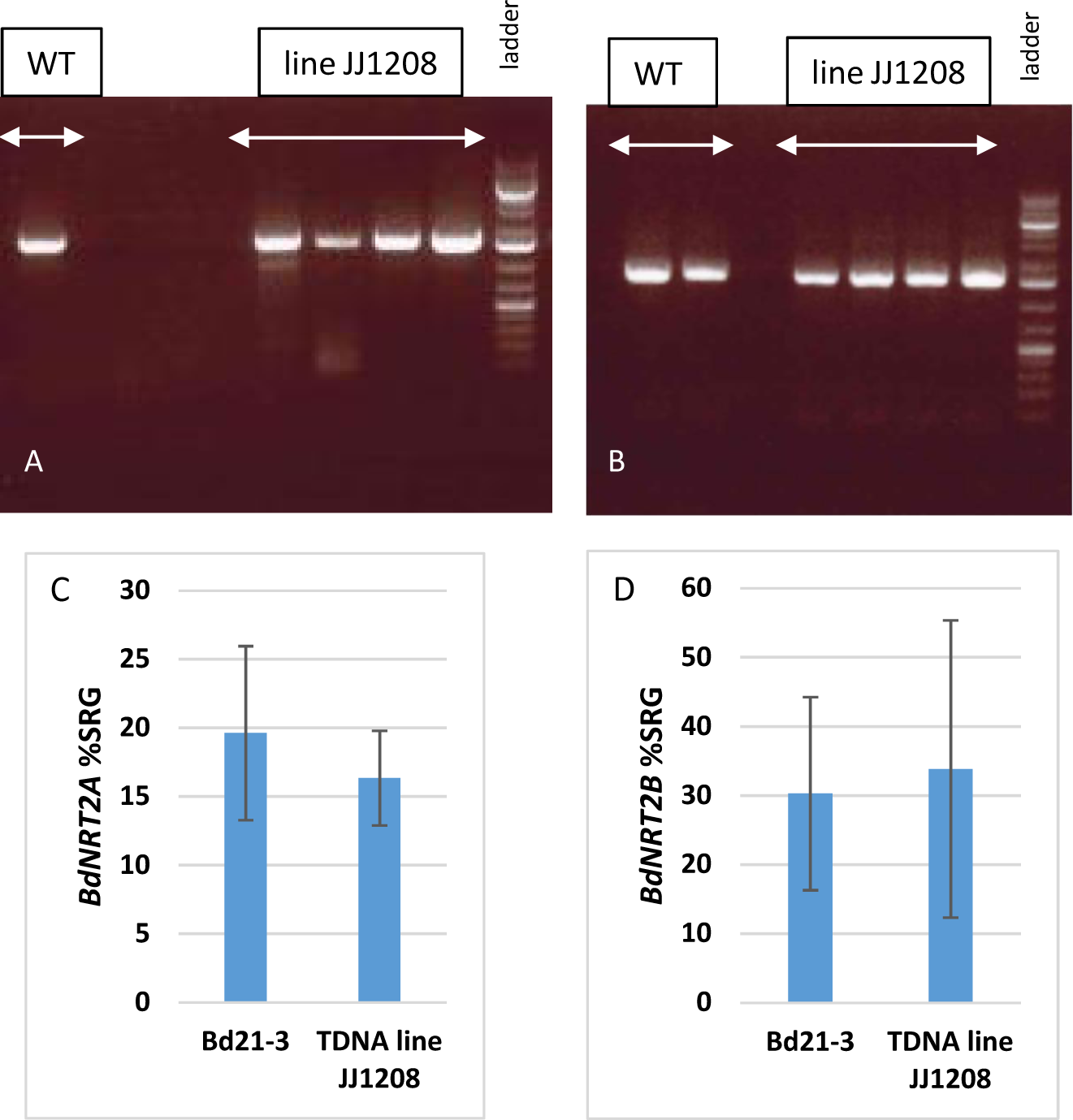
Absence of a T-DNA insertion in the *BdNRT2A* gene in line JJ1208 (obtained from the JGI Brachypodium collection T-DNA mutants in the Bd21-3 accession) demonstrated by the quantification of *BdNRT2A* and *BdNRT2B* gene expression in roots of plants homozygous for the resistance to hygromycin. S1A-B. Gene expressions were measured by RT-PCR (using specific primers (*BdNRT2AspecifF* and *70R2* for *BdNRT2A*) (S1A) and (*BdNRT2BspecifF* and *70R2* for *BdNRT2B*) (S1B) (annealing temperature 54°C, 35 cycles of amplification). Products of amplification (1.5kb) were separated by electrophoresis on 1% agar gel, and GeneRuler^TM^ 1kb plus ladder (Fermentas) was used for the size quantification. Four independant plants are shown for the T-DNA line JJ1208. S1C-D. Gene expressions were quantified by qRT-PCR using specific primers (see list of primers) for *BdNRT2A* (S1C) and *BdNRT2B* (S1D) and were normalized to the level of a synthetic reference gene (SRG). *BdUBI10* and *BdSAMDC* were used to compose the SRG. Values are means ± SD of 4 biological replicates.

**Supplementary data S2 :**
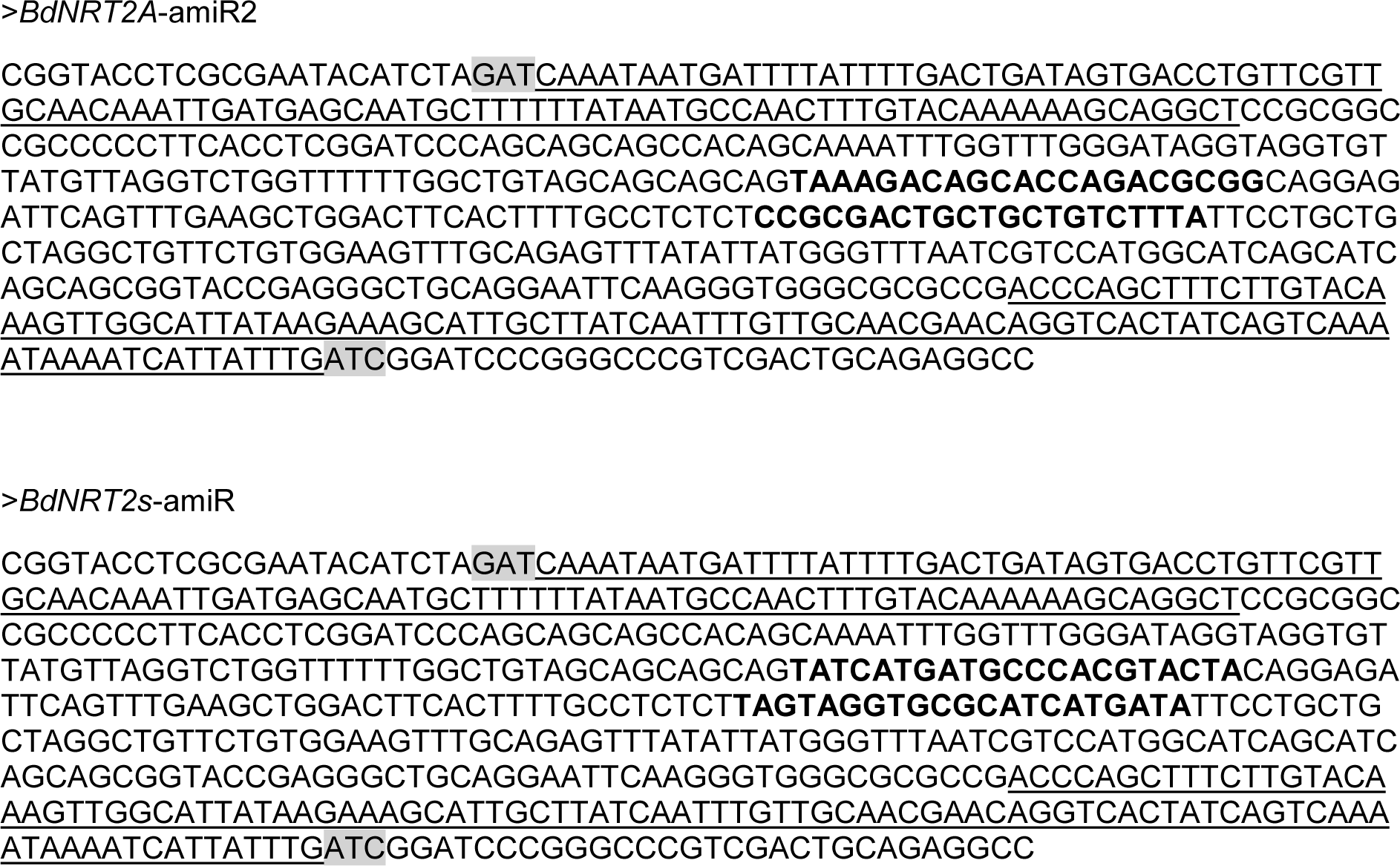
Sequences of amiRNA constructs in pUC57-Kan. Target-specific sequences are in bold; *attL1* and *attL2* sequences for Gateway LR reaction are underlined; remains of *EcoRV* sites used to clone the fragments in pUC57-Kan are highlighted in grey. These constructs allowed the generation of two amiRNA mutant lines (*amiRj2* with *BdNRT2s*-amiR and *amiRn3* with *BdNRT2s*-amiR) as described in the results.

**Supplementary data S3 :**
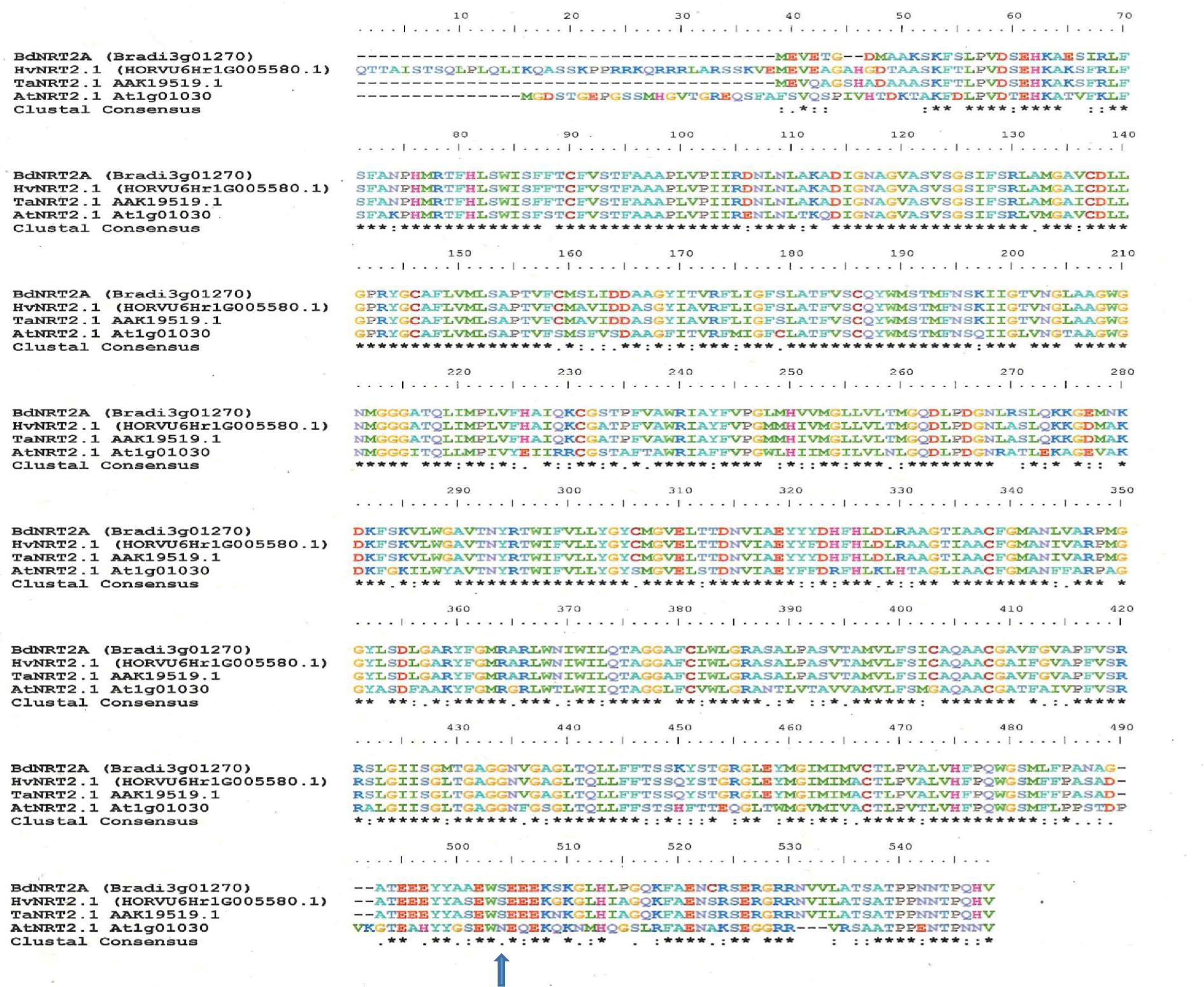
Alignement of amino acid sequences of BdNR2A, HvNR2.1, TaNRT2.1 and AtNRT2.1. The Ser residue not conserved in dicotyledons is indicated with an arrow. Sequence alignments were performed using ClustalW Multiple alignment application available in Bioedit sequence alignment editor.

## References

Bogard M, Allard V, Brancourt-Hulmel M, Heumez E, MacHet JM, Jeuffroy MH, Gate P, Martre P, Le Gouis J. 2010. Deviation from the grain protein concentration-grain yield negative relationship is highly correlated to post-anthesis N uptake in winter wheat. Journal of Experimental Botany 61: 4303–4312.

Cai C, Wang JY, Zhu YG, Shen QR, Li B, Tong YP, Li ZS. 2008. Gene structure and expression of the high-affinity nitrate transport system in rice roots. Journal of Integrative Plant Biology 50: 443–451.

Cleaver OB, Patterson KD, Krieg PA, 1996. Overexpression of the tinman-related genes XNkx-2.5 and XNkx-2.3 in Xenopus embryos results in myocardial hyperplasia. Dev. Camb. Engl. 122, 3549– 3556.

Chen J, Zhang Y, Tan Y, Zhang M, Zhu L, Xu G, Fan X. 2016. Agronomic nitrogen-use efficiency of rice can be increased by driving OsNRT2.1 expression with the OsNAR2.1 promoter. Plant Biotechnology Journal 14: 1705–1715.

Chopin F, Orsel M, Dorbe MF, Chardon F, Truong HN, Miller AJ, Krapp A, Daniel-Vedele F. 2007. The Arabidopsis ATNRT2.7 nitrate transporter controlsnitrate content in seeds. Plant Cell.;19(5):1590–602.

Citovsky V, Lee LY, Vyas S, Glick E, Chen MH, Vainstein A, Gafni Y, Gelvin SB, Tzfira T. 2006. Subcellular Localization of Interacting Proteins by Bimolecular Fluorescence Complementation in Planta J. Mol. Biol., 362, pp. 1120–1131.

Clough, S.J. and Bent, AF. 1998 Floral dip : a simplified method for Agrobacterium mediated transformation of Arabidopsis thaliana. Plant J. 16, 735–743.

Dalmais M, Antelme S, Ho-Yue-Kuang S, Wang Y, Darracq O, d’Yvoire MB, Cézard L, Légée F, Blondet E, Oria N, et al. 2013. A TILLING Platform for Functional Genomics in Brachypodium distachyon. PLoS ONE 8.

David LC, Berquin P, Kanno Y, Mitsunori S, Daniel-Vedele F and Ferrario-Méry S. 2016 N availability modulates the role of NPF3.1, a gibberellin transporter, in GA-mediated phenotypes in *Arabidopsis*. Planta 244, 1315–1328.

David LC, Girin T, Fleurisson E, Phommabouth E, Mahfoudhi A, Citerne S, Berquin P, Daniel-Vedele F, Krapp A, Ferrario-Méry S. 2019. Developmental and physiological responses of Brachypodium distachyon to fluctuating nitrogen availability. Scientific Reports 9. 3824.

Dechorgnat J, Patrit O, Krapp A, Fagard M, Daniel-Vedele F. 2012 Characterization of the Nrt2.6 Gene in Arabidopsis haliana: A Link with Plant Response to Biotic and Abiotic Stress. PLoS ONE 7(8): e42491. doi:10.1371/journal.pone.0042491

Engelsberger WR, Schulze WX 2012 Nitrate and ammonium lead to distinct global dynamic phosphorylation patterns when resupplied to nitrogen-starved Arabidopsis seedlings. Plant J 69: 978– 995

Faure-Rabasse S, Deunff E Le, Macduff JH, Laine P, Ourry A. 2002. Effects of nitrate pulses on BnNRT1 and BnNRT2 genes : mRNA levels and nitrate influx rates in relation to the duration of N deprivation in Brassica napus L . Journal of Experimental Botany 53: 1711–1721.

Feng H, Yan M, Fan X, Li B, Shen Q, Miller AJ, Xu G. 2011. Spatial expression and regulation of rice high-affinity nitrate transporters by nitrogen and carbon status. 62: 2319–2332.

Filleur S, Daniel-Vedele F. 1999. Expression analysis of a high-affinity nitrate transporter isolated from Arabidopsis thaliana by differential display. Planta 207: 461–496.

Filleur S, Dorbe MF, Cerezo M, Orsel M, Granier F, Gojon A, Daniel-Vedele F. 2001. An Arabidopsis T-DNA mutant affected in Nrt2 genes is impaired in nitrate uptake. FEBS Letters 489: 220– 224.

Girin T, David LC, Chardin C, Sibout R, Krapp A, Ferrario-Méry S, Daniel-Vedele F. 2014. Brachypodium: A promising hub between model species and cereals. Journal of Experimental Botany 65: 5683–5686.

Himmelbach A, Zierold U, Hensel G, Riechen J, Douchkov D, Schweizer P, Kumlehn J. 2007. A set of modular binary vectors for transformation of cereals. Plant physiology 145, 1192–200.

Hong S-Y, Seo P J, Yang M-S, Xiang F. and Park C-M. 2008 Exploring valid reference genes for gene expression studies in Brachypodium distachyon by real-time PCR. BMC Plant Biol. 8, 112.

Ishikawa S, Ito Y, Sato Y, Fukaya Y, Takahashi M, Morikawa H, Ohtake N, Ohyama T, Sueyoshi K. 2009. Two-component high-affinity nitrate transport system in barley: Membrane localization, protein expression in roots and a direct protein-protein interaction. Plant Biotechnology 26: 197–205.

Jacquot A, Chaput V, Mauries A, Li Z, Tillard P, Fizames C, Bonillo P, Bellegarde F, Laugier E, Santoni V, et al. 2020. NRT2.1 C-terminus phosphorylation prevents root high affinity nitrate uptake activity in Arabidopsis thaliana. New Phytologist 228: 1038–1054.

Jacquot A, Li Z, Gojon A, Schulze W, Lejay L. 2017. Post-translational regulation of nitrogen transporters in plants and microorganisms. Journal of Experimental Botany 68: 2567–2580.

Kawachi T, Sunaga Y, Ebato M, Hatanaka T, Harada H. 2006. Repression of nitrate uptake by replacement of Asp105 by asparagine in AtNRT3.1 in Arabidopsis thaliana L. Plant and Cell Physiology 47: 1437–1441.

Kiba T, Feria-Bourrellier A-B, Lafouge F, Lezhneva L, Boutet-Mercey S, Orsel M, Bréhaut V, Miller A, Daniel-Vedele F, Sakakibara H, et al. 2012. The Arabidopsis Nitrate Transporter NRT2.4 Plays a Double Role in Roots and Shoots of Nitrogen-Starved Plants . The Plant Cell 24: 245–258.

Kotur Z, Mackenzie N, Ramesh S, Tyerman S D, Kaiser B N and Glass A D M 2012 Nitrate transport capacity of the Arabidopsis thaliana NRT2 family members and their interactions with AtNAR2.1 New Phytol 194: 724–731

Kotur Z, Unkles SE, Glass ADM. 2017. Comparisons of the Arabidopsis thaliana High-af fi nity Nitrate Transporter Complex AtNRT2 . 1 / AtNAR2 . 1 and the Aspergillus nidulans AnNRTA : structure function considerations. 64: 3–4.

Laugier E, Bouguyon E, Mauriès A, Tillard P, Gojon A, and Lejay L. 2012. Regulation of High-Affinity Nitrate Uptake in Roots of Arabidopsis Depends Predominantly on Posttranscriptional Control of the. Plant Physiology 158: 1067–1078.

Lejay L, Tillard P, Lepetit M, Olive FD, Filleur S, Daniel-Vedele F, Gojon A. 1999. Molecular and functional regulation of two NO3/-uptake systems by N- and C-status of Arabidopsis plants. Plant Journal 18: 509–519.

Lezhneva L, Kiba T, Feria-Bourrellier AB, Lafouge F, Boutet-Mercey S, Zoufan P, Sakakibara H, Daniel-Vedele F, Krapp A. 2014. The Arabidopsis nitrate transporter NRT2.5 plays a role in nitrate acquisition and remobilization in nitrogen-starved plants. Plant Journal 80: 230–241.

Li W, Wang Y, Okamoto M, Crawford NM, Siddiqi MY, Glass ADM. 2007. Dissection of the AtNRT2.1 : AtNRT2.2 Inducible High-Affinity Nitrate Transporter Gene Cluster . Plant Physiology 143: 425–433.

Liu X, Huang D, Tao J, Miller AJ, Fan X, Xu G. 2014. Identification and functional assay of the interaction motifs in the partner protein OsNAR2.1 of the two-component system for high-affinity nitrate transport. New Phytologist 204: 74–80.

Menz J, Li Z, Schulze WX, Ludewig U (2016) Early nitrogen deprivation responses in Arabidopsis roots reveal distinct differences on transcriptome and (phospho-) proteome levels between nitrate and ammonium nutrition. Plant J 88: 717–734

Miranda, K. M., Espey, M. G. & Wink, D. A. 2001 A Rapid, Simple Spectrophotometric Method for Simultaneous Detection of Nitrate and Nitrite. Nitric Oxide 5, 62–71.

Nakamura Y, Umemiya Y, Masuda K, Inoue H, Fukumoto M. 2007. Molecular cloning and expression analysis of cDNAs encoding a putative Nrt2 nitrate transporter from peach. Tree Physiology 27: 503–510.

Nazoa P, Vidmar JJ, Tranbarger TJ, Mouline K, Damiani I, Tillard P, Zhuo D, Glass ADM, Touraine B. 2003. Regulation of the nitrate transporter gene AtNRT2.1 in Arabidopsis thaliana: Responses to nitrate, amino acids and developmental stage. Plant Molecular Biology 52: 689–703.

Okamoto M, Kumar A, Li, W, Wang Y, Siddiqi MY, Crawford NM, Glass ADM. 2006. High-Affinity Nitrate Transport in Roots of Arabidopsis Depends on Expression of the NAR2-Like Gene AtNRT3.1. PLANT Physiol. 140, 1036–1046.

Okamoto M, Vidmar JJ, Glass ADM. 2003. Regulation of NRT1 and NRT2 gene families of Arabidopsis thaliana: responses to nitrate provision. Plant and Cell Physiology 44: 304–317.

Orsel M, Chopin F, Leleu O, Smith SJ, Krapp A, Daniel-Vedele F, Miller AJ. 2006. Characterization of a Two-Component High-Affinity Nitrate Uptake System in Arabidopsis. Physiology and Protein-Protein Interaction. Plant Physiology 142: 1304–1317.

Orsel M, Chopin F, Leleu O, Smith SJ, Krapp A, Daniel-Vedele F, Miller AJ. 2007. Nitrate signaling and the two component high affinity uptake system in Arabidopsis. Plant Signal Behav. 2(4):260–2.

Orsel M, Eulenburg K, Krapp A, Daniel-Vedele F. 2004. Disruption of the nitrate transporter genes AtNRT2.1 and AtNRT2.2 restricts growth at low external nitrate concentration. Planta 219: 714–721.

Ono F, Frommer W B and von Wiren N. 2000. Coordinated diurnal regulation of low- and high-affinity nitrate transporters in tomato. Plant Biology 2: 17–23.

Quesada A, Krapp A, Trueman L J, Daniel-Vedele F, Fernandez E, Forde B G, Caboche M. 1997. PCR-identification of a Nicotiana plumbaginifolia cDNA homologous to the high-affinity nitrate transporters of the crnA family. Plant Molecular Biology 34: 265–274.

Tong Y, Zhou JJ, Li Z, Miller AJ. 2005. A two-component high-affinity nitrate uptake system in barley. Plant Journal 41: 442–450.

Vandesompele J, De Preter K, Pattyn F, Poppe B, Van Roy N, De Paepe A, Speleman F. 2002. Accurate normalization of real-time quantitative RT-PCR data by geometric averaging of multiple internal control genes. Genome Biology 3, RESEARCH0034.

Vidmar JJ, Zhuo D, Siddiqi MY, Glass ADM. 2000. Isolation and Characterization of HvNRT2.3 and HvNRT2.4, cDNAs Encoding High-Affinity Nitrate Transporters from Roots of Barley . Plant Physiology 122: 783–792.

Vogel J and Hill T 2008 High-efficiency Agrobacterium-mediated transformation of Brachypodium distachyon inbred line Bd21-3. Plant Cell Rep, 27(3):471–478.

Wang Y-Y, Cheng Y-H, Chen K-E, Tsay Y-F. 2018. Nitrate Transport, Signaling, and Use Efficiency. Annual Review of Plant Biology 69.

Wang J, Huner N and Tian L 2019 Identification and molecular characterization of the Brachypodium distachyon NRT2 family, with a major role of BdNRT2.1. Physiologia Plantarum 165: 498–510.

Warthmann N, Chen H, Ossowski S, Weigel D, Hervé P. 2008. Highly specific gene silencing by artificial miRNAs in rice. PloS one 3, e1829.

Wirth J, Chopin F, Santoni V, Viennois G, Tillard P, Krapp A, Lejay L, Daniel-Vedele F, Gojon A. 2007. Regulation of root nitrate uptake at the NRT2.1 protein level in Arabidopsis thaliana. Journal of Biological Chemistry 282: 23541–23552.

Yan M, Fan X, Feng H, Miller AJ, Shen Q, Xu G. 2011. Rice OsNAR2.1 interacts with OsNRT2.1, OsNRT2.2 and OsNRT2.3a nitrate transporters to provide uptake over high and low concentration ranges. Plant, Cell and Environment 34: 1360–1372.

Yin LP, Li P, Wen B, Taylor D, Berry JO. 2007. Characterization and expression of a high-affinity nitrate system transporter gene (TaNRT2.1) from wheat roots, and its evolutionary relationship to other NTR2 genes. Plant Science 172: 621–631.

Yong Z, Kotur Z, Glass ADM. 2010. Characterization of an intact two-component high-affinity nitrate transporter from Arabidopsis roots. Plant Journal 63: 739–748.

Zhai Z, Jung H, Vatamaniuk OK. 2009. Isolation of protoplasts from tissues of 14-day-old seedlings of Arabidopsis thaliana. JoVE. 30. http://www.jove.com/details.php?id=1149, doi: 10.3791/1149

Zhuo D, Okamoto M, Vidmar JJ, Glass ADM. 1999. Regulation of a putative high-affinity nitrate transporter (Nrt2;1At) in roots of Arabidopsis thaliana. Plant Journal 17: 563–568.

Zhou JJ, Theodoulou FL, Muldin I, Ingemarsson B, Miller AJ. 1998. Cloning and functional characterization of a Brassica napus transporter that is able to transport nitrate and histidine. J. Biol. Chem. 273, 12017–12023.

Zou X, Liu MY, Wu WH, Wang Y. 2020. Phosphorylation at Ser28 stabilizes the Arabidopsis nitrate transporter NRT2.1 in response to nitrate limitation. Journal of Integrative Plant Biology 62: 865–876.

